# Adaptive integration of self-motion and goals in posterior parietal cortex

**DOI:** 10.1101/2020.12.19.423589

**Authors:** Andrew S. Alexander, Janet C. Tung, G. William Chapman, Laura E. Shelley, Michael E. Hasselmo, Douglas A. Nitz

## Abstract

Animals engage in a variety of navigational behaviors that require different regimes of behavioral control. In the wild, rats readily switch between foraging and more complex behaviors such as chase, wherein they pursue other rats or small prey. These tasks require vastly different tracking of multiple behaviorally-significant variables including self-motion state. It is unknown whether changes in navigational context flexibly modulate the encoding of these variables. To explore this possibility, we compared self-motion processing in the multisensory posterior parietal cortex while rats performed alternating blocks of free foraging and visual target pursuit. Animals performed the pursuit task and demonstrated predictive processing by anticipating target trajectories and intercepting them. Relative to free exploration, pursuit sessions yielded greater proportions of parietal cortex neurons with reliable sensitivity to self-motion. Multiplicative gain modulation was observed during pursuit which increased the dynamic range of tuning and led to enhanced decoding accuracy of self-motion state. We found that self-motion sensitivity in parietal cortex was history-dependent regardless of behavioral context but that the temporal window of self-motion tracking was extended during target pursuit. Finally, many self-motion sensitive neurons conjunctively tracked the position of the visual target relative to the animal in egocentric coordinates, thus providing a potential coding mechanism for the observed gain changes to self-motion signals. We conclude that posterior parietal cortex dynamically integrates behaviorally-relevant information in response to ongoing task demands.

## Introduction

Goal-directed navigation takes many complex forms across species. Many animals, including rats and mice, exhibit pursuit behaviors in social, sexual, and predatory contexts (Calhoun, 1963; Eisenberg and Leyhausen, 1972; Kurtz and Adler, 1973). Pursuit navigation requires continuous adjustment of movement plans as a function of past and current movement states and the position of the moving goal relative to the animal (i.e. in egocentric space). In primates, efficiency of chase behaviors is enhanced by predictive movements based on the hypothesized trajectory of the target through space (Barnes et al., 2000; Yoo et al., 2020), though this has yet to be reported in murine animals.

The demanding nature of pursuit behavior requires greater precision of sensorimotor processing than other forms of navigation, such as those observed during foraging for food sources. Movement is known to modulate sensory processing (Niell and Stryker, 2010; Vinck et al., 2015; Bouvier et al., 2020; Guitchounts et al., 2020), but the influence of behaviorally-relevant sensory information on movement coding has received little attention in freely moving rodents. Namely, it is unclear whether the context of navigation (e.g. pursuit or foraging) can flexibly modulate movement processing in support of ongoing behavioral demands. Here, we sought to explore the effect of navigational context on sensorimotor processing by examining pursuit navigation in rats.

In rodents, chase behaviors require the dorsal striatum, superior colliculus (SC), and zona incerta which mediate goal detection, pursuit initiation, and orienting behaviors required for continuous pursuit (Schiller and Stryker, 1972; Cooper et al., 1998; Hoy et al., 2016, 2019; Shang et al., 2019; Zhao et al., 2019; Kim et al., 2019). Although much has been revealed about these regions and predatory chase, little is known about cortical involvement. Multisensory cues that define the context of predatory behavior (e.g. the sight, sound, or smell of prey) are potentially combined in association cortices such as posterior parietal cortex (PPC) which possesses reciprocal connectivity to sensorimotor cortical regions and projects to the SC and dorsal striatum (Reep et al., 1994; Wilber et al., 2015; Hovde et al., 2019; Gilissen et al., 2020). PPC damage produces hemispatial neglect, which manifests as an inability to detect or orient to stimuli present in space contralateral to the site of lesion (Bisiach and Luzzatti, 1978; Behrmann et al., 1997). Thus, PPC functioning is critical for spatial awareness and attentional processes necessary to chase a moving goal.

Relevant to motor coordination during chase, PPC neurons exhibit sensitivity to self-motion (e.g. linear and angular speed), posture, and visual target position and movement direction in egocentric coordinates (Kawano et al., 1980; Chen et al., 1994; Whitlock et al., 2012; Rancz et al., 2015; Wilber et al., 2017; Andersen and Mountcastle, 1983; Wilber et al., 2014; Sasaki et al., 2020; Mimica et al., 2018). PPC also computes multimodal abstractions of space and movement commands in manners important for pursuit behavior. PPC ensembles are sensitive to the shape of trajectories through space, animal position within a route, task phase in both spatial and non-spatial domains, and the integration of information over time for decision making (Nitz, 2006, 2012; Harvey et al., 2012; Goard et al., 2016; Minderer et al., 2019; Scott et al., 2017; Hwang et al., 2017; Akrami et al., 2018; Krumin et al., 2018; Morcos and Harvey, 2016). Further, PPC is hypothesized to support coordinate system transformations which may subserve online sensorimotor coordination necessary for target chasing (Save and Poucet, 2000; Angelaki and Cullen, 2008; Bicanski and Burgess, 2018). Collectively, these signals may drive initiation of predation behaviors and orienting motor commands in PPC target structures.

To explore PPC dynamics during pursuit behavior we recorded neural activity extracellularly while rats chased and intercepted a floor projected visual stimulus to receive reward. We tracked the animal’s position and the location of the visual target simultaneously and were thus able to examine whether PPC ensembles were responsive to multiple variables relevant to pursuit behavior including self-motion and the egocentric position of the target. In separate experimental sessions on the same day, animals freely foraged for food in the same arena which enabled a comparison of self-motion processing as a function of navigational context. Rats readily learned the task and exhibited predictive behaviors consistent with those observed in primates. We report that PPC coding was flexibly modulated as a function of navigation behavior, indicating that the region may play a significant role in integrating multisensory cues and engaging navigation circuitry in response to behavioral context.

## Results

### Rats perform a target pursuit task and exhibit shortcutting behavior

Rats (n = 6) performed a target chasing task wherein they pursued and attempted to intercept an experimenter controlled light target moving along the surface of an open arena in experimenter-generated pseudorandom trajectories (RTs). On each trial, the animal engaged in pursuit behavior until it subjectively caught the light target (**Figure 1a, sFigure 1a, sVideos 1-3,**). Following interception, the light target switched off, a reward was delivered to a random position within the arena, and the light target reappeared hovering on the reward to guide retrieval. After consumption, the next trial began at varying environmental positions dependent on the animals heading and location. During each recording session, rats averaged 34 pursuits (n = 132 sessions, median = 33.5, Interquartile range (IQR) = 24 – 41.5) of a median 3.95 seconds in duration (IQR = 2.74 – 5.42s).

**Figure 1.**
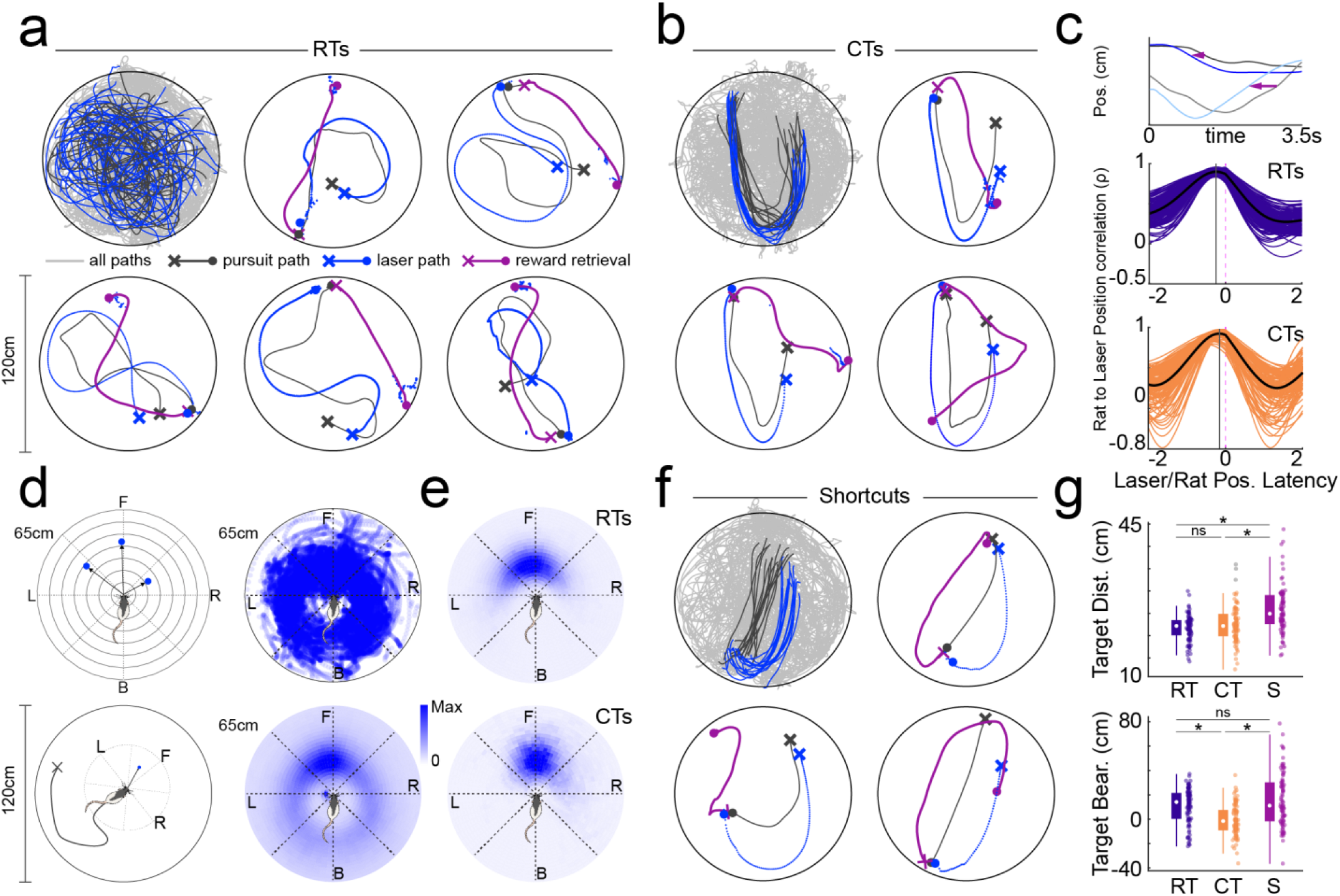
Rats pursue visual targets and exhibit spatial shortcuts on known trajectories. **a.** Rats chase a floor-projected visual target moving in pseudorandom trajectories (RTs). Top left, all paths throughout an example target pursuit session (light gray), all pseudorandom pursuit paths (dark gray), and all pseudorandom paths of the visual target (blue). Remaining plots depict 5 example pursuits with trial starting location (X), approximate location of target interception (circle), and path to reward retrieval (purple) marked. Rat trajectories are depicted in light gray and laser trajectories are in blue. **b.** Top left, all paths throughout an example target pursuit session (light gray), all pursuits along the characteristic trajectory (CTs; dark gray), and all characteristic paths of the visual target (blue). Remaining plots depict 3 example characteristic trajectories marked as in **a**. **c.** Quantification of temporal relationship between the animal and visual target. Top plot, animal (x, dark gray; y, light gray) and laser position (x, blue; y, light blue) across time on an example characteristic trajectory trial. Arrows illustrate temporal lag between rat and visual target position. Middle plot, correlation between rat and target position as a function of temporal shift of rat position relative to the target during pseudorandom trajectories (RTs). Black line is mean of all trials and light gray lines depict individual sessions. Bottom plot, same as above but for characteristic trajectories (CTs). Position correlation curves are right shifted during CTs relative to RTs indicating that rat and target position was more temporally proximal during CTs. Vertical gray lines depict latency with peak correlation for each trial type. **d.** Mapping of visual target relative to the rat in egocentric coordinates. Top left, scheme for examining egocentric relationship between visual target and rat. For each position frame, the animal is positioned at the origin of the polar axes with current heading oriented upward and the egocentric bearing and distance to the visual target are mapped to the angular and radial axes, respectively. Bottom left, schematic depicting example egocentric position of target relative to the rat independent of allocentric position or heading in the 120cm diameter circular environment. Top right, all visual target positions relative to the animal during mobility within an example pursuit session. Bottom right, mean target occupancy in egocentric coordinates across all animals and pursuit sessions (n = 132). **e.** Top, mean target occupancy in egocentric coordinates for all pseudorandom trajectories across all animals and sessions. Bottom, same as above but for all characteristic trajectories. **f.** Same as in **a** and **b**, but for characteristic trajectory trials in which the animal executed a spatial shortcut. **g.** Median distance and bearing to the visual target for pseudorandom (RT), characteristic (CT), and shortcut (S) trajectories.

To explore the effect of navigational uncertainty on animal behavior we randomly inserted “characteristic trajectories (CT)” of the moving visual target amongst the pseudorandom trajectories (**Figure 1b, sFigure 1b, sVideos 4-6,** n = 13 CTs per session, IQR = 10 – 17). Each instance of a CT started and finished at approximately the same locations within the allocentric reference frame (as defined by visible distal cues), and possessed a reliable shape that involved moving outbound from the initiation point to the opposite side of the arena, performing a U-turn, and returning inbound to a location displaced approximately 25cm from the start point (**Figure 1b**).

The presence of CTs introduced regularity in an otherwise stochastic navigational environment for the animal. We questioned whether the animals could learn the shape of the CT path through path integration despite its immersion among random paths and that its outline was never physically or visually present at any given moment in time. Thus, we hypothesized that after learning the existence of such a path, the animal would be able to predict the trajectory of the moving target during CTs and more closely track the stimulus when compared to RTs. Indeed, temporal shifts of rat position relative to visual target position revealed that animals were approximately 250ms (IQR = 200 – 300ms) behind the location of the visual stimulus during random pursuits, but only 150ms (IQR = 100ms – 200ms) behind during CTs (**Figure 1c,** gray vertical line, mean temporal lag yielding highest correlation between rat and target position; n = 132, Wilcoxon rank sum test, z = 6.19, p = 5.94×10^−10^).

To further characterize pursuit behavior we next examined the distance and angle of the light stimulus relative to the animal in egocentric coordinates, irrespective of allocentric position within the environment or current heading (**Figure 1d**). Throughout the entirety of the pursuit block, which included both target pursuits and subsequent reward retrievals, the visual target occupied the full range of egocentric bearings and distances relative to the animal (**Figure 1d**). During actual pursuits, target position was highly correlated between random and characteristic trajectories and primarily localized to a 90° range centered along the nasal axis of the animal (**Figure 1e**). Distance to the visual target was slightly greater and more variable during CTs when compared to RTs (RT median distance = 20.4cm, IQR = 18.2 – 22.4; CT median = 21.7cm, IQR = 19.1 – 23.3, Wilcoxon rank sum test, z = −2.36, p = 0.02; RT median distance range = 9.5cm, IQR = 8.39 – 11.3, CT median distance range = 11.5cm, IQR = 9.45 – 14.1, z = −3.73, p = 1.86×10^−4^). Egocentric bearing to the target was unchanged between trial types (RT median = −2.41° (CCW), IQR = −17.4 – 11.6; CT median = 4.53 (CW), IQR = −4.13 – 13.8, Wilcoxon rank sum test, z = 1.93, p = 0.05), but the range of target angles was substantially reduced during CTs (RT median bearing range = 50.4°, IQR = 44.1 – 56.9; CT median bearing range = 28.4°, IQR = 23.3 – 38.2, Wilcoxon rank sum test, z = 8.63, p = 5.96×10^−18^).

The increased maintenance of a consistent angular position relative to the visual cue on CTs indicated that rats had internalized the sequence of actions, headings, and positions that collectively composed these known paths. In support of this interpretation, rats demonstrated knowledge of the structure of CTs by exhibiting shortcutting behavior (**Figure 1f,** median shortcuts of CTs per session 20%, IQR = 6.7 – 44.9%; n = 3 shortcuts trials per session, IQR = 1 – 5.5). Shortcuts primarily involved interception of the visual target at the apex of the CT trajectory (**Figure 1f**, **sVideos 7-9**), but took many different forms (**sFigure 1c**). Interception of the visual target as a behavioral strategy required the animal to navigate through the region bordered by outbound and inbound segments of the CT. Accordingly, shortcuts could be distinguished from non-shortcut trials in two ways. First, rat distance to target increased on shortcut trials prior to interception (**Figure 1g,** distance shortcuts = 24.9cm, IQR = 22.6 – 29.1 cm; CT = 22.1cm, IQR = 19.9 – 24.9cm; RT = 22.1cm, IQR = 20.1 – 23.5cm; Kruskal-Wallis test w/ Tukey-Kramer, n = 91, χ^2^ = 39.5, p =2.6×10^−9^). Secondly, egocentric bearing to the target was increased on shortcuts when compared to full CTs (**Figure 1g,** median bearing shortcuts = 11.0°, IQR = −1.7 – 30.2°; CT = −1.7°, IQR = −9.1 – 7.5°; RT = 13.9°, IQR = 0.3 – 21.5°cm; Kruskal-Wallis test w/ Tukey-Kramer, n = 91, χ^2^ = 39.8, p =2.3×10^−9^). Overall, the shortcutting behavior indicated that the rat could recognize the structure of the outbound component of the stable path trials and predict the future trajectory of the visual target.

These observations demonstrate that rats can be trained to pursue moving targets that are purely visual in nature. This ecologically-inspired paradigm mimics capture and chase behaviors that rats exhibit in the wild. To efficiently perform this task, animals must actively integrate sensorimotor and spatial information of target trajectories and exhibit adaptive self-motion behavior. The execution of shortcuts indicate that rodents can learn, and therefore predict, these target trajectories to guide subsequent action sequences in a spatiotemporally insightful manner.

### Nonlinear self-motion correlates of PPC cells

Chase behavior requires online integration of visual, motor, and spatial information. The significance of these signals, and their interactions, likely varies between target pursuit and more self-guided behavior such as free foraging. We hypothesized that navigational context would modulate self-motion related neural activity within posterior parietal cortex (PPC), which possesses dense reciprocal connectivity with visual and sensorimotor processing areas.

We recorded 367 neurons from posterior parietal cortex (PPC, n = 5 rats, n = 132 sessions) while animals performed the target chasing paradigm (**sFigure 2**). To examine modulation of self-motion correlates as a function of task context, we paired a subset of target pursuit sessions with preceding and/or succeeding free foraging sessions within the same arena and no visual target present (n = 113 sessions, n = 302 PPC neurons). Pursuit blocks were longer in temporal duration than foraging in order to yield adequate numbers of pursuits (pursuit block median length = 22.7 minutes, IQR = 21.1 – 25.1; free explore median length = 12.3 minutes, IQR = 10.9 – 15.7), but animals only engaged in target chasing behavior for a brief amount of time within this period (median time pursuing target = 3.02 minutes, IQR = 2.32 – 3.92).

**Figure 2.**
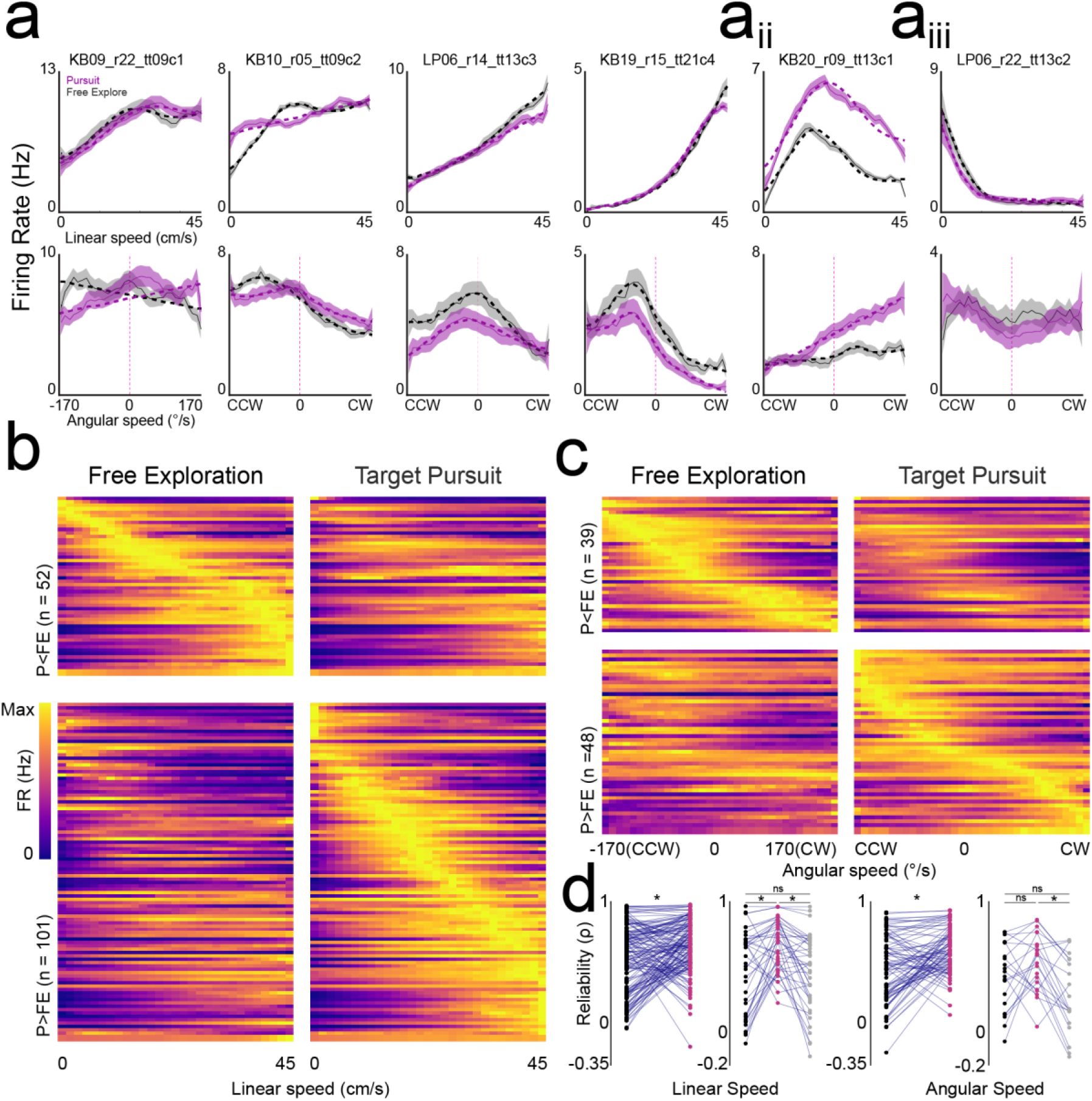
Nonlinear self-motion correlates of PPC are modulated by navigational demands. **a.** Linear (top) and angular (bottom) speed tuning curves for 6 PPC neurons (columns, mean ± s.e.). Dashed lines are best model fit and are shown only for tuning curves that had significant modulation; aii, An example neuron with Gaussian-like nonlinear linear speed tuning; aiii, An example neuron with robust firing during immobility. **b.** Linear speed tuning curves for both free exploration and target pursuit epochs for all PPC neurons with significant modulation. Top plots, linear speed tuning curves for neurons that had greater mean activation during free exploration (left column), sorted by peak linear speed bin in free explore. Bottom plots, linear speed tuning curves for neurons that had greater mean activation during target pursuit (right column), sorted by peak linear speed bin in target pursuit. **c.** Same as in **b**, but for neurons with significant angular speed tuning. **d.** Left plots, comparison of reliability in linear speed tuning curves for neurons recorded during free explore (black dots) and pursuit (pink dots) or for the subset of neurons recorded in pursuit (pink dots) and in two bounding free explore sessions (black and gray dots). Reliability is systematically increased during target pursuit. Right plots, same as left plots but for neurons with significant angular speed tuning.

As a consequence of the more restricted task structure, rats exhibited lower median linear speeds during the full pursuit session when compared to free exploration (**sFigure 3a**, n = 113 sessions, speed free exploration = 21.5 cm/s, IQR = 17.5 – 24.6 cm/s; full pursuit = 13.9 cm/s, IQR = 11.4 – 17.0 cm/s). However, rats were indeed engaged in the task as linear speeds during actual target pursuits were greater than those observed in free exploration (**sFigure 3a,** median linear speed in pursuit = 47.6 cm/s, IQR = 44.8 – 51.7, Kruskal-Wallis test w/ Tukey-Kramer, n = 113, χ^2^ = 257.9, p =1.01×10^−56^). In contrast, absolute angular velocity (AV) was highly similar between the full pursuit block and free foraging, but was reduced during actual pursuit epochs indicating that trajectories dictated by the path of the visual target were considerably more smooth with less abrupt angular deviations (**sFigure 3b**, n = 113 sessions, average absolute AV free exploration = 74.9 deg/s, IQR = 70.1 – 78.8 deg/s; full pursuit = 78.9 deg/s, IQR = 74.9 – 83.8 deg/s; actual pursuits = 59.0 deg/s, IQR = 53.6 – 71.6 deg/s; Kruskal-Wallis test w/ Tukey-Kramer, n = 113, χ^2^ = 86.8, p =1.41×10^−19^). Autocorrelations of linear and angular speed further supported that there was greater regularity (e.g. smoothness) in self-motion behavior during target chasing than in free exploration (**sFigure 3c-d).**

**Figure 3.**
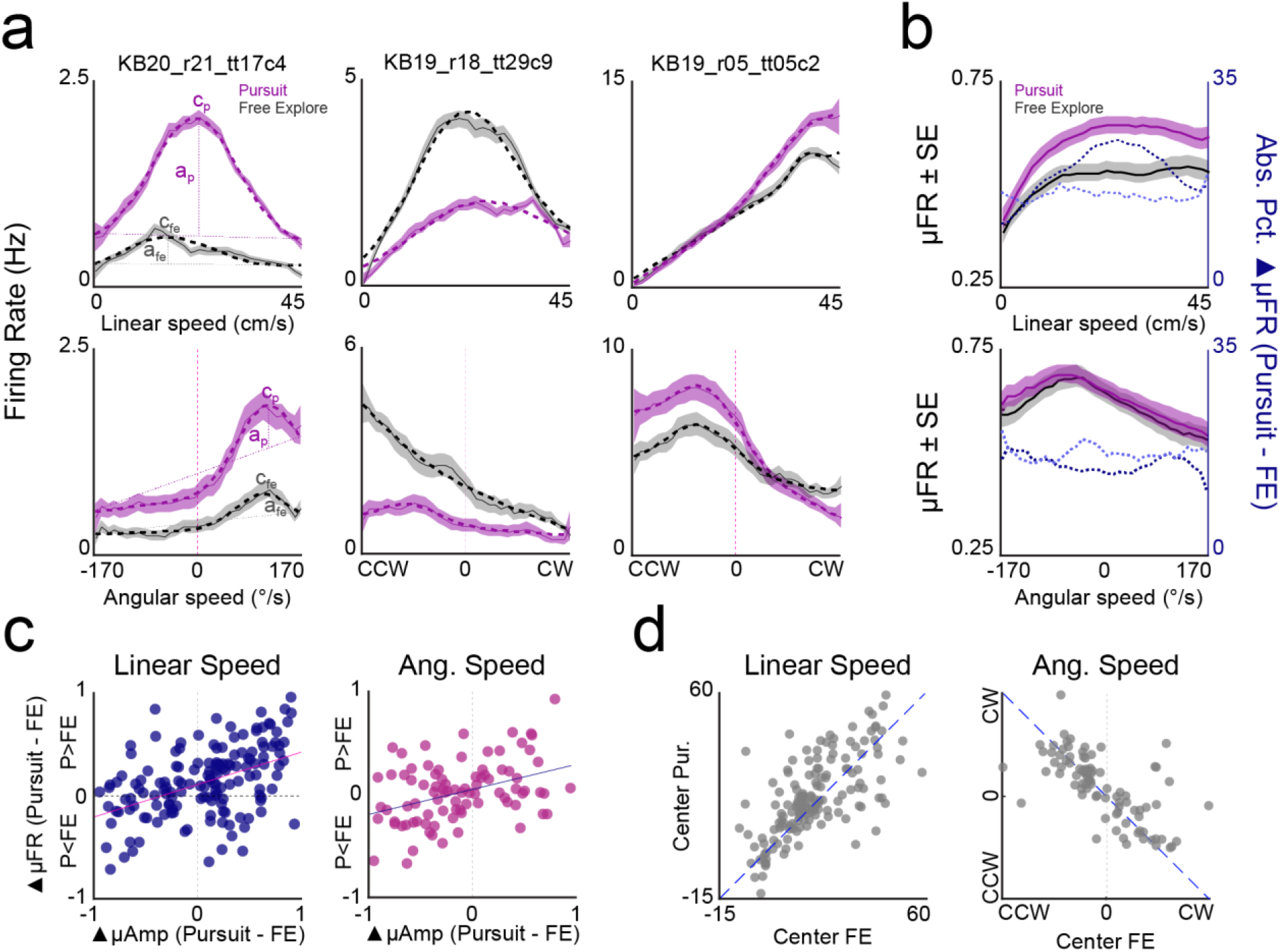
Gain modulation of self-motion tuning as a function of navigational demand. **a.** Three example PPC neurons with rate differences between free exploration and pursuit that are concentrated at specific linear (top row) or angular (bottom row) speeds. Left plot schematizes parameters of Gaussian-modified linear fits. Sloped dashed lines indicate linear regression. Self-motion receptive fields are fit by an additive Gaussian function. The center (c) and amplitude (a) of the Gaussian map the peak of the self-motion receptive field and its magnitude above the linear fit, respectively. **b.** Shaded line plots depict mean population tuning for linear (top) and angular (bottom) speed sensitive neurons for pursuit (pink) and free exploration sessions (gray). Dashed lines depict mean percent difference in firing rate as a function of speed (dark blue, PPC neurons with greater activation in pursuit than foraging [P>FE]; light blue, PPC neurons with greater activation in foraging [P<FE]). **c.** Peak normalized differences in receptive field amplitude between pursuit and free explore (▲μAmp) versus difference in mean firing rate (▲μFR) for PPC neurons with significant linear and/or angular speed correlates. Positive values indicate greater amplitude/mean rate in pursuit. Positive correlation between amplitude and mean rate differences for both linear (left) and angular (right) sensitive populations suggests neurons exhibit both additive and multiplicative gain modulation wherein mean and within receptive field firing rates are increased in the neurons “preferred” behavioral session. **d.** Scatterplots depict the position of linear and angular receptive fields for pursuit and free exploration. Preferred linear speeds occurred at higher speeds for pursuit than free explore (left plot). Preferred angular speeds were preserved between behaviors (right plot).

As we sought to compare neural correlates of free exploration and target chasing sessions despite systematic variation in self-motion between epochs, we next calculated linear and angular velocity tuning curves for each task for all PPC neurons (**Figure 2a**) as well as for randomizations wherein the spike train was shifted relative to behavior 1000 times. All analyses of neural data during the pursuit session were conducted on the full block including pursuit trials, reward consumption, and intertrial intervals. In all cases, tuning curves were constructed based on matched sampling of linear and angular velocities. We fit true and randomized tuning curves with a uniform distribution, a linear model, and a Gaussian-modified linear model (GML) which featured parameters for both linear regression and Gaussian fitting. Neurons with significant linear or GML model fit (relative to the uniform fit) and significant within session reliability (relative to the distribution of within session reliability values calculated from randomizations) were deemed to possess linear and/or angular speed tuning.

56.6% (n = 171/302) of PPC neurons met this criterion for either linear or angular speed tuning during either target pursuit or free exploration (**Figure 2b-c**). Linear speed tuning was more prominent in PPC (**Figure 2b,** 50.6%, n = 153/302) than angular speed related activation (**Figure 2c,** 28.8%, n = 87/302), and most angular tuned neurons were simultaneously sensitive to linear speed (79.3% n=69/87). 51.0% of all putative principal neurons were influenced by self-motion (45.3% linear velocity modulated, n = 111/245 principal cells, 25.3% angular velocity modulated, n = 62/245 principal cells, n = 62/87 (71.3%) of all angular velocity modulated including interneurons). Nearly all putative interneurons that were sensitive to movement (80.7%, n = 46/57) were modulated by linear speed (91.3% n = 42/46) while fewer were modulated by angular velocity (54.3%, n = 25/46).

For most cells, the relationship between firing rate and self-motion appeared nonlinear. To quantify this observation, we compared model fit between the linear model and the GML model for all tuned neurons (see methods). Tuning curves of most linear speed sensitive cells showed significant improvement to model fit when Gaussian terms were included in addition to linear terms, indicating high incidence of nonlinear relationships to linear speed within the region (**Figure 2a, dashed lines**; 75.8%, n = 116/153, *F*-test linear vs. linear + Gaussian, *F*(3,24) = [3.01– 4480.2], *p* = [0 – 0.049]). Many nonlinear speed relationships resembled previously reported saturating speed tuning (**Figure 2a,** Hinman et al., 2016) while others had distinct firing rate peaks at specific linear speeds (**Figure 2aii**). This latter group included a subset of neurons with activity peaks during immobility (**Figure 2aiii**). Most tuning curves of angular velocity sensitive PPC cells were also best fit by Gaussian-modified linear models (87.4%, n = 76/87, *F*-test linear vs. linear + Gaussian, *F*(3,29) = [2.96 – 130.43], *p* = [0 – 0.049]). Many angular speed tuning curves had pronounced Gaussian characteristics reflective of receptive fields spanning restricted ranges of angular movements rather than linear functions transitioning between clockwise and counterclockwise movements (**Figure 2a, 2c**).

### Self-motion correlates are recruited by navigational context and more reliable during chasing

We next questioned whether navigational context, as defined by target pursuit or free exploration, was reflected in the self-motion related activity of PPC. Consistent with flexible recruitment of ensembles during increased navigational demands, the number of neurons with self-motion related activation increased during pursuit behavior when compared to free exploration (FE linear speed, n = 96/302, 31.8%; Pursuit linear speed, n = 126/302, 41.7%; FE angular speed, n = 45/302, 14.9%; Pursuit angular speed, n = 72/302, 23.8%). Individual neurons exhibited similar firing as a function of self-motion between pursuit and free exploration as assessed via Spearman’s correlation of their tuning curves, even if the neuron only reached criterion for detection in one condition (**Figure 2b-c,** Spearman’s *ρ* FE vs. Pursuit, median *ρ* linear speed = 0.60, IQR = 0.37-0.79; median *ρ* angular speed = 0.57, IQR = 0.38-0.74).

Based on these results, we hypothesized that the increase in significantly tuned self-motion sensitive neurons during pursuit could arise from 1) increased within session reliability of tuning curves during pursuit, a criterion for detection of significant tuning (see methods) and/or 2) increased signal-to-noise ratio (i.e. dynamic range of firing rate modulation), arising from adaptive nonlinear dynamics between self-motion and neuronal activation. The latter might be expected to yield greater proportions of significantly non-uniform tuning profiles.

To examine the first possibility, we assessed the stability of self-motion sensitivity within free exploration and target pursuit. For each neuron we computed the correlation of tuning curves independently constructed from non-overlapping odd and even minutes within each behavioral epoch. Reliability of both linear and angular speed correlates were significantly greater during pursuit when compared to free explore (**Figure 2d,** linear speed, n = 153, median reliability (ρ) FE = 0.57, IQR = 0.25 – 0.73, median reliability Pursuit = 0.65, IQR = 0.48 – 0.80, Wilcoxon rank sum, z = − 3.38, p = 0.0007; angular speed, n = 87, median reliability FE = 0.44, IQR = 0.18 – 0.67, median reliability Pursuit = 0.565, IQR = 0.44 – 0.72, z = −3.56, p =0.0004). Increased reliability of linear speed tuning was systematically related to pursuit behavior, as reliability values returned to baseline levels in a second free exploration session following the pursuit epoch (Kruskal-Wallis test on linear speed reliability FE1 vs Pursuit vs FE2, w/ post-hoc Tukey Kramer, n = 40, χ^2^ = 11.5, p =0.003; Kruskal-Wallis test on angular speed reliability FE1 vs Pursuit vs FE2, w/ post-hoc Tukey Kramer, n = 21, χ^2^ = 7.61, p = 0.02). These findings indicate that PPC dynamically increases the precision of self-motion related correlates as an adaptation to increased navigational demands during target chasing.

### PPC self-motion correlates exhibit two forms of gain modulation as a function of navigational context

We next investigated the second hypothesis by looking for the presence of gain modulation in self-motion related firing rate relationships. We began by examining mean firing rate differences in self-motion tuning taken from free exploration and pursuit. Sub-populations of PPC neurons exhibited increases or decreases in mean overall response as a function of self-motion and behavioral epoch (**sFigure 4a**). **Figures 2b** and **2c** show all linear and angular speed tuning curves for both free exploration and pursuit blocks, split into subsets with increased (P>FE) or decreased (P<FE) firing rate during pursuit and sorted by position of peak self-motion tuning within their “preferred” navigational epoch.

**Figure 4.**
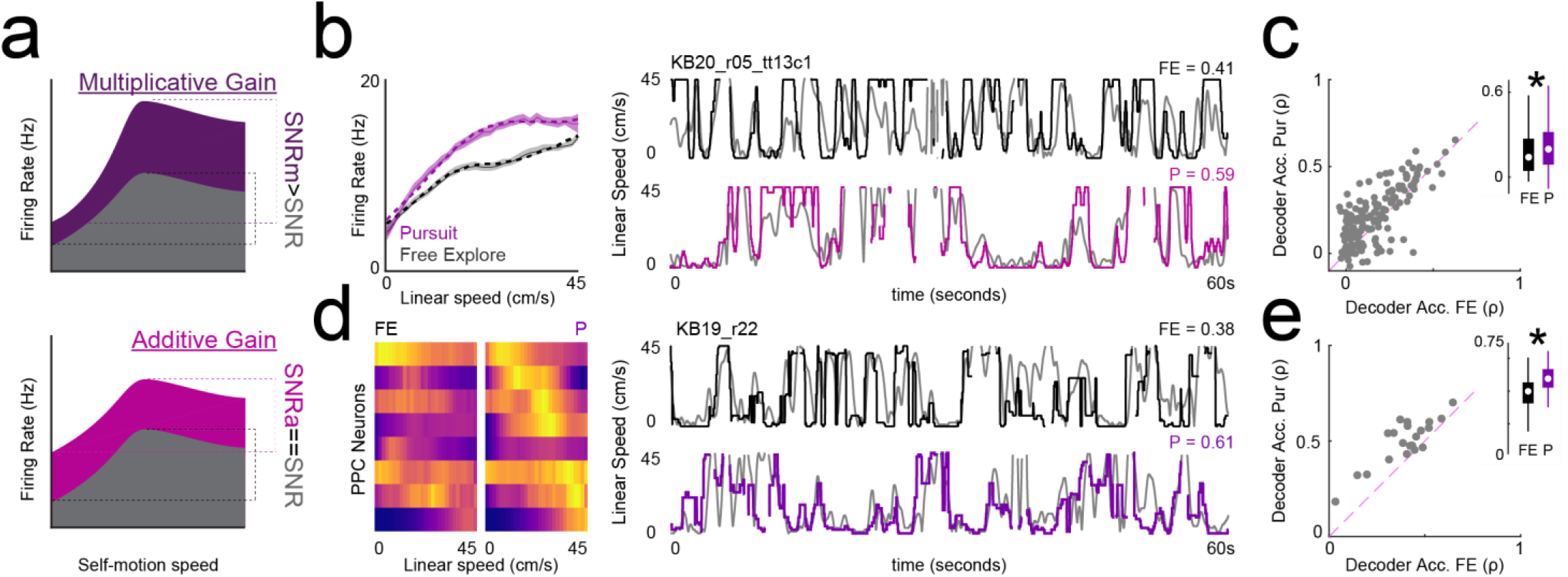
Multiplicative gain modulation produces enhanced dynamic range and decoding of self-motion correlates. **a.** Schematic of multiplicative (top) and additive (bottom) gain modulation on self-motion tuning curves. Gray curve depicts hypothetical relationship between self-motion and firing rate for a baseline session. Colored curves depict hypothetical relationship between self-motion and firing rate for a session in which either multiplicative or additive gain modulation manifests. In the case of additive gain, the signal-to-noise ratio (SNR) is equivalent between the modulated session (SNRa==SNR) while multiplicative gain produces an enhanced SNR for the modulated session (SNRm>SNR). **b.** Left, example linear speed sensitive neuron with multiplicative gain modulation. Right, decoding of linear speed using spiking activity of the same neuron in FE (top) and pursuit (bottom). Decoding accuracy was assessed by correlating the predicted speed (black/pink) with the true speed (gray) and is indicated above each plot. **c.** Linear speed decoder accuracy (Spearman’s rho) is greater for pursuit than FE for a majority of neurons. Inset, median and IQR of decoder accuracy for FE (black) and pursuit (purple). **d.** Left plot, linearized linear speed tuning curves for 8 simultaneously recorded PPC neurons in FE and P. Rows correspond to the linear speed tuning curves of the same neurons recorded in both conditions and the colormap indicates low (purple) to maximum firing (yellow), peak normalized across all FE and pursuit sessions. Right, corresponding decoder output using this ensemble for FE (top) and pursuit (bottom). Decoder accuracy is indicated above each plot. **e.** Decoder accuracy as a function of session, presented as in d, but for ensemble decoding shows enhanced linear speed decoding in pursuit.

Significant proportions of linear speed modulated cells had greater mean firing rates during the pursuit epoch than the foraging session (Binomial test for proportion of linear speed cells with greater activity in pursuit vs. FE, n = 153, n P>FE = 101, p = 9.17×10^−5^; angular speed cells, n = 87, n P>FE = 48, p = 0.39). Linear speed sensitive neurons with increased firing during pursuit (P>FE) returned to baseline firing during a second free exploration session supporting the interpretation that the increased rate was directly related to pursuit behavior (**sFigure 4b**). Systematic modulation of mean rate as a function of behavioral context was not preserved across all self-motion tuned neurons. Neurons that decreased their firing during pursuit did not have consistent rate relationships to task. Instead, PPC neurons composing the P<FE subset showed greater similarity in mean firing rate for neighboring sessions, suggesting that the firing rate changes could arise from ensemble drift or sustained state changes following task engagement during target chasing (**sFigure 4b**).

Mean rate differences reflect additive gain modulation on self-motion responsivity but could ignore more dynamic changes to firing rate between navigational tasks. For example, self-motion tuning differences between pursuit and FE could be restricted to specific ranges of movement speeds consistent with multiplicative gain modulation (**Figure 3a, Figure 4a**). We next explored self-motion related firing rate changes at a finer resolution by 1) investigating the distribution of firing rate differences between behaviors as a function of self-motion state and 2) examining parameters of self-motion tuning curves.

Pursuit and free foraging tuning curves diverged as a function of self-motion speed and revealed different patterns for linear and angular speed tuned neurons (**Figure 3b**). Specifically, linear speed tuning appeared to exhibit increased differences in firing rate between pursuit and free foraging when the animal was running at higher speeds. Angular speed tuning exhibited task-specific modulation of firing rate magnitude that was more uniformly distributed across angular speeds (**Figure 3b**). The profile of rate differences between pursuit and FE was associated with: 1) the distribution of peaks in self-motion related activation (**Figure 2b-c**) and, 2) the profile of mean tuning across the full population of neurons (**Figure 3b)**. This indicated that, in addition to mean rate changes between tasks, rate differences may also be concentrated to self-motion states associated with peak activation for each neuron. **Figure 3a** depicts 3 example PPC neurons that illustrate this effect.

To examine within receptive field firing rate modulation between sessions, we compared parameters of Gaussian-modified linear model (GML) fits to self-motion tuning curves. As the GML model implements linear regression with additive 1-D Gaussian parameters to capture nonlinearities, the amplitude parameter (a) reflects additional activation above a linear fit, and thus models within receptive field firing magnitude (as schematized in **Figure 3a**). PPC neurons exhibited significant amplitude alterations to self-motion sensitivity that were consistent with multiplicative gain modulation (**Figure 3c**, **sFigure 5a-b**). As was the case with mean rate changes between behaviors, amplitude gain alterations were, in most but not all cases, modulated in a systematic fashion relative to task context (**sFigure 5c**). Within-receptive field amplitude differences were correlated with alterations of mean rate between pursuit and free exploration, indicating that many neurons were subject to both additive and multiplicative gain modulation of their self-motion related activation between navigational contexts (**Figure 3c, Figure 4a**).

**Figure 5.**
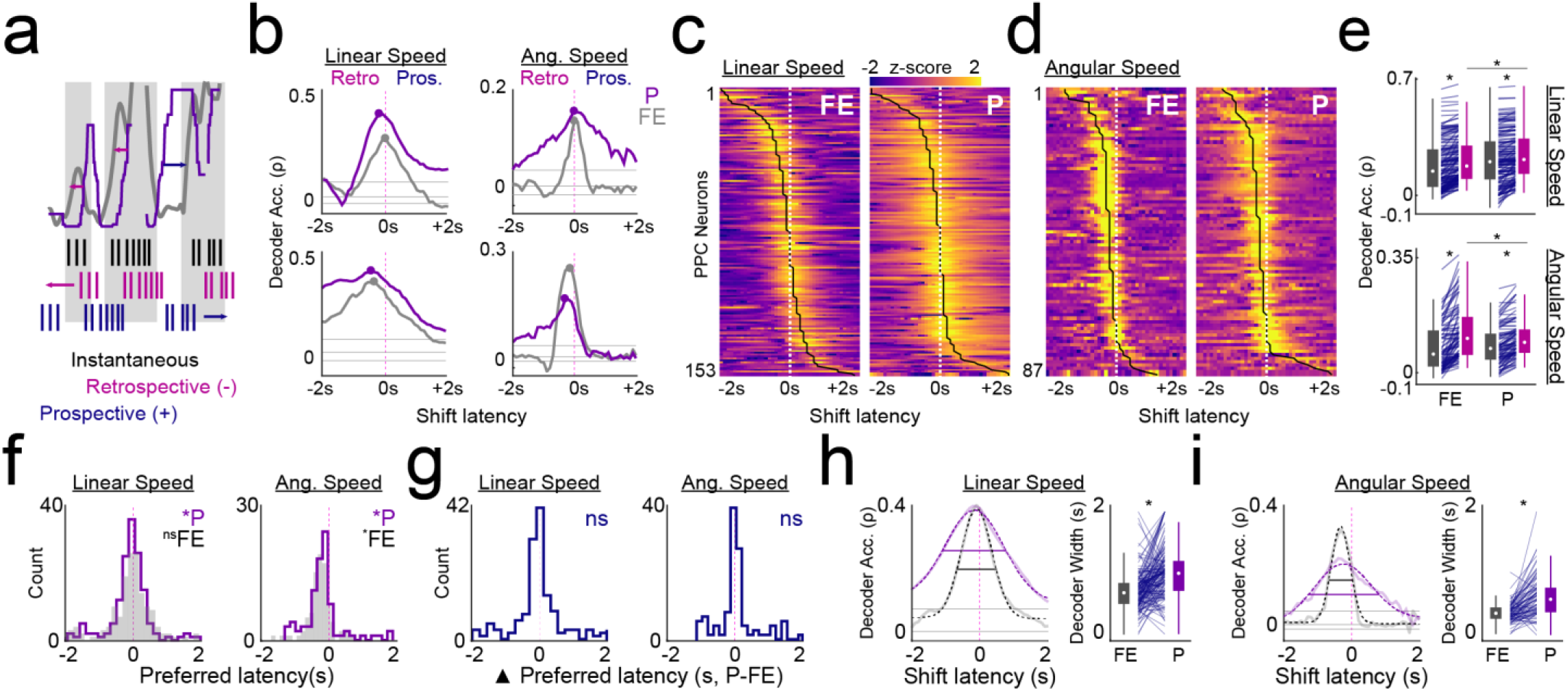
History-dependent spiking correlates are informative about self-motion state over extended temporal windows. **a.** Schematic depicting temporal relationship between spiking activity, self-motion, and decoder output. On top, gray and purple lines depict real and predicted linear speed, respectively. Gray shading indicates times at which the rats speed is changing. Colored arrows depict direction of spike train temporal shift that could yield increased decoder accuracy. Below, hypothetical spike trains for a single neuron either instantaneously (black), retrospectively (pink), or prospectively (blue) sensitive to speed. **b.** Decoder accuracy (Spearman’s ρ) as a function of shift of spike train in time relative to behavior for 4 PPC neurons in both free exploration (gray) and pursuit conditions (purple). Left column, 2 PPC neurons sensitive to linear speed. Right column, 2 PPC neurons sensitive to angular speed. Increased decoder accuracy for negative shifts of the spike train reflect history-dependence (retrospective) of spiking relative to self-motion, while increased accuracy for positive shifts of the spike train indicate an anticipatory (prospective) relationship to self-motion. Gray horizontal lines indicate the 99th, 50th, and 1st percentiles of decoding accuracy following multiple shifts of the spike train at latencies greater than 2s. Dots indicate spike train temporal shift yielding peak decoding accuracy (i.e. preferred latency). **c.** Decoding accuracy latency curves for all linear speed sensitive neurons, sorted by latency with peak decoding within free explore (left) and pursuit (right). **d.** Same as in **c**, but for all angular speed sensitive neurons. **e.** Comparison between decoding accuracy at instantaneous (gray) and preferred (pink, value at shift latency yielding maximum decoding accuracy) relationships between spiking and self-motion for free explore (FE) and pursuit session (P). In both behavioral epochs, decoding accuracy was significantly greater at non-instantaneous shift latencies for linear (top) and angular (bottom) speed sensitive neurons. **f.** Distribution of preferred shift latencies for linear and angular speed neurons in pursuit and free exploration is skewed to negative shifts indicating retrospective relationships between spiking and self-motion. **g.** Histogram showing non-significant differences in preferred latency between pursuit and free exploration for linear and angular speed neurons. **h.** Left, example decoding accuracy latency curve for a linear speed sensitive cell in pursuit and free exploration with corresponding model fits (dashed lines). Colored horizontal bars are plotted at approximately 50% of the peak to visualize width differences between the sessions. Right, decoder width is significantly greater in pursuit than free exploration for linear speed sensitive neurons indicating that prediction from spiking is accurate for extended temporal windows in pursuit. **i.** Same as in **h**, but for the temporal window of angular speed decoding.

In pursuit, receptive field amplitude was increased for preferred linear speeds and decreased for preferred angular speeds (**sFigure 5a**). These changes were reminiscent of animal behavior during active target chasing wherein rats typically exhibited higher linear speeds and lower angular speeds when compared to free exploration (**sFigure 3**). In this respect, we recall that our tuning curve construction controls for differences in self-motion across contexts by matching the sampling of rates at each speed. In parallel with the pattern of changes to receptive field amplitude, the center parameter (c) of model fits (as schematized in **Figure 3a**), corresponding to the location of the Gaussian component of the fits, shifted to higher linear speeds during pursuit when compared to free explore (**Figure 3d,** Wilcoxon sign rank test for zero median, n = 153, z = 2.47, p = 0.0134). Angular speed preferences were unchanged between pursuit and free explore (Wilcoxon sign rank test for zero median Pursuit – FE, n = 87, z = −1.38, p = 0.17). Thus, both the gain and profile of self-motion tuning curves were modulated to match animal behavior, indicating that self-motion correlates in PPC are highly adaptive to behavioral demands.

### Decoding of self-motion state is modulated by navigational context

The presence of multiplicative gain modulation, rather than purely additive gain, suggested that the signal-to-noise ratio (SNR) for self-motion coding was altered as a function of task and overall enhanced during pursuit (**Figure 4a**). If changes to SNR are meaningful, they should yield corresponding changes to the resolution of self-motion decoding in downstream readers. To test this hypothesis, we trained a cross-validated (ten-fold) maximum correlation coefficient classifier on the relationship between instantaneous firing rate (computed by convolving the spike train with a Gaussian filter with 200ms standard deviation, see methods) and linear and angular speed for each neuron independently (**Figure 4b, sFigure 6**). We then predicted the linear and angular speed from corresponding instantaneous firing rates at each position sample omitted from the training set. Classifier accuracy was quantified by correlating the predicted linear or angular speed with the true value as exemplified in **Figure 4b**.

**Figure 6.**
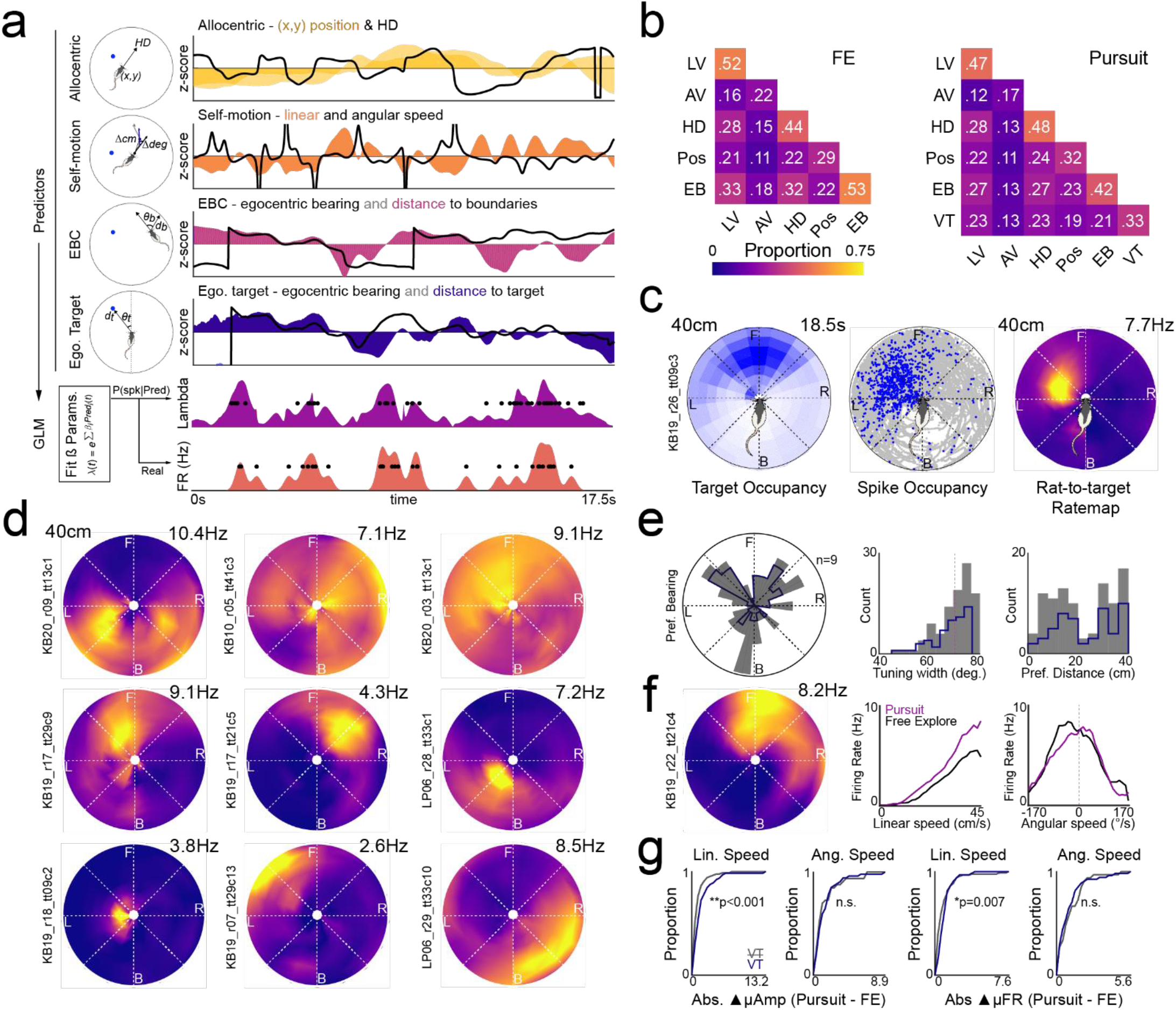
PPC neurons track self-motion and the egocentric position of the visual target. **a.** Schematic of generalized linear modeling framework for the assessment of sensitivity to multiple spatial and behavioral variables simultaneously. Left column, schematic illustrations depicting different predictor classes. Right column, 17.5 seconds of z-score normalized values for the different predictor classes. Directional predictors (HD, angular speed, egocentric bearing to boundary, egocentric bearing to target) depicted in black lines, all other predictors presented in shaded colors. Bottom plots, GLM fits ß weight parameters to each predictor and generates a probability of spiking for each timestamp (lambda) which can be compared to the real spike train. **b.** Proportion of all PPC neurons (n = 302) sensitive to each predictor class (along the diagonal) as well as all pairwise combinations of predictor classes (off diagonal) for free exploration (left, FE) and pursuit (right). LV, linear velocity; AV, angular velocity; HD, head direction; Pos, allocentric position; EB, egocentric boundary; VT, visual target. **c.** Rat-to-target ratemaps. Left plot, heatmap depicts egocentric occupancy of target relative to the rat throughout the session as shown in **Figure 1d** (white low occupancy, blue high occupancy). Middle, trajectory plot depicting all positions of the target relative to the rat in gray and all positions in which a single neuron spiked in blue. Right, rat-to-target ratemap constructed after binning and normalizing the spike occupancies from the middle plot by the total target occupancy. This neuron is active when the visual target is to the animal’s front left. **d.** 9 example rat-to-target ratemaps for PPC neurons with significant sensitivity to the egocentric position of the visual target as assessed via the GLM. Top row, 3 neurons with broad bearing selectivity and limited target distance information. Bottom two rows, 6 neurons with more restricted target position receptive fields possessing both bearing and distance components. **e.** Properties of egocentric target receptive fields. Left, polar distribution depicting preferred bearing of all PPC neurons with significant tuning to the target in gray. In dark blue, preferred bearing for PPC neurons with reliable bearing across non-overlapping halves of the target chasing session. Middle plot, distribution of width of egocentric bearing tuning. Right, distribution of preferred distances to visual target. All colors as in left plot. **f.** An example PPC neuron with sensitivity to the egocentric position of the visual target and self-motion simultaneously. Left plot, rat-to-target ratemap. Middle plot, linear speed tuning curves in pursuit (purple) and free explore (black). Right, angular speed tuning curves for both sessions. This neuron exhibited gain modulation of linear speed coding during pursuit. **g.** Left two plots, cumulative density functions depict the absolute difference in receptive field amplitude (i.e. multiplicative gain) between pursuit and free exploration for linear and angular speed sensitive neurons conjunctively sensitive to target position (VT, purple) or not (VT, gray). Significant rightward shift of VT curve indicates neurons with target sensitivity exhibited greater gain modulation. Right two plots, same as left plots, but for absolute difference in mean rate (i.e. additive gain) between self-motion sensitive neurons with or without VT sensitivity. Linear speed sensitive neurons that also coded target position had significantly greater additive and multiplicative gain than the subset of neurons that were not sensitive to the target.

Consistent with the hypothesized relationship between SNR alterations and self-motion processing, linear speed decoding was significantly more accurate during pursuit for most PPC neurons with linear speed sensitivity (**Figure 4c**, Binomial test for decoding accuracy of linear speed pursuit vs. FE, n = 153, n P>FE = 103, p = 2.20×10^−5^; Wilcoxon rank sum test, median accuracy (*ρ*) Pursuit = 0.20, IQR = 0.1 – 0.32; *ρ* FE = 0.14, IQR = 0.05 – 0.28, z = 2.35, p = 0.02). There was no significant bias in angular speed decoding accuracy in pursuit versus free foraging (**sFigure 6**, Binomial test, pursuit vs. FE, n = 87, n P>FE = 49, p = 0.28; Wilcoxon rank sum test, z = 1.16, p = 0.25).

Similar self-motion decoding results were observed when simultaneously recorded ensembles (n ≥ 5 neurons, n = 22 sessions) were utilized (**Figure 4d,e**, **sFigure6**, Wilcoxon rank sum test, median ensemble decoding accuracy (*ρ*) of linear speed Pursuit = 0.5, IQR = 0.45 – 0.57; median *ρ* FE = 0.42, IQR = 0.34 – 0.48, z = 2.48, p = 0.013; angular speed *ρ* pursuit = 0.12, IQR = 0.08 – 0.15; angular speed *ρ* FE = 0.13, IQR = 0.09 – 0.17, z = 1.19, p = 0.24). The relationship between accuracy of linear and angular speed decoding and pursuit versus free exploration again matched the task-dependent behavior of the animal, further supporting adaptive engagement of cortical ensembles for tracking navigationally-relevant variables as a function of task demands.

### Self-motion decoding is accurate for extended temporal windows during pursuit behavior

Spiking activity can exhibit temporally latent relationships to ongoing behavior with individual neurons exhibiting prospective or retrospective activation in relation to movement or other variables. Qualitative examination of decoder output revealed temporal offsets between self-motion predictions and true linear or angular speed values (**Figure 5a**). Accordingly, we questioned whether systematic shifts in the temporal latency between spike trains and behavior would alter decoder accuracy. We shifted each neuron’s instantaneous firing rate vector 2 seconds backwards or forwards in time in 100ms increments relative to behavior and decoded linear and angular speed using the same decoding approach. Importantly, if decoder accuracy increased during backwards shifts, this implied that activation of the neuron was correlated with the history of movement (i.e. retrospectively), while enhanced predictions following forward spike train shifts suggested that activation of the neuron was prospectively related to the locomotor state (**Figure 5a**).

**Figure 5b** depicts the profile of decoding accuracy for four example PPC neurons following iterative shifts of the spike train relative to behavior (left column, two example linear speed sensitive neurons; right column, angular speed sensitive neurons). For most neurons, the preferred latency yielding maximal decoder accuracy was observed at non-instantaneous spike train lags (**Figure 5c-d**). Self-motion decoding accuracy at preferred latencies was significantly greater than the zero-lag instantaneous decoding for both free exploration and target chasing sessions (**Figure 5e**, Wilcoxon sign rank test, linear speed, n = 153, free explore, z = −10.04, p = 9.8×10^−24^, pursuit, z = −9.78, p = 1.39×10^−22^; angular speed, n = 87, free explore, z = −7.77, p = 7.85×10^−15^, pursuit, z = −7.67, p =1.68×10^−14^). However, maximum overall prediction accuracy, across all possible spike train offsets relative to linear speed, was still significantly greater during target chasing despite decoding enhancements observed in free explore (i.e. decoding at the preferred latency during free exploration did not surpass decoding accuracy at the preferred latency in target pursuit; Wilcoxon sign rank test, linear speed, n = 153, z = −3.75, p = 0.0001). The opposite pattern was observed for angular speed decoding accuracy which was greater during free exploration across all possible latencies (Wilcoxon sign rank test, linear speed, n = 153, z = −3.75, p = 0.0001; angular speed, n = 87, z = 2.91, p = 0.004).

At the population level, preferred latencies of linear speed decoding appeared significantly skewed in the retrospective direction (**Figure 5c-d).** To examine this statistically, we fit the profile of decoding accuracy latency curves with the Gaussian-modified linear model and compared the center parameters between the two behavioral tasks. Preferred latencies for linear speed decoding across the population were biased toward history-dependence in both foraging and target pursuit yet only reached significance from the instantaneous latency in pursuit (**Figure 5f**, Wilcoxon sign rank for zero median, linear speed, n = 153, median free explore = −10.9ms, IQR = −313.5 – 210.7ms, z = − 1.14, p = 0.26; median pursuit = −57.3ms, IQR = −280.3 – 181.4ms, z = −2.12, p = 0.03). Similar, yet even more retrospective, distributions of preferred latencies were observed in angular speed decoding (**Figure 5f**, Wilcoxon sign rank for zero median, angular speed, n = 87, median free explore = −252.8ms, IQR = −371.7 – −106.2ms, z = −6.08, p = 1.19×10^−9^; median pursuit = −189.4ms, IQR = − 411.4 – −66.1ms, z = −4.75, p = 2.02×10^−6^). Overall, there were no significant changes in preferred latency for individual neurons between pursuit and free exploration, indicating that although parietal cortex neurons are more responsive to the history of self-motion, this preference is rigid and not task dependent (**Figure 5g**, Wilcoxon sign rank for zero median difference in preferred latencies, linear speed, z = −1.17, p = 0.24; angular speed, z = 1.34, p = 0.18).

While there were no differences between pursuit and free exploration in the preferred time lag in the relationship between spiking and self-motion, examination of the temporal profile of decoding accuracy suggested that PPC neurons may exhibit extended temporal windows of movement integration as a function of task context (**Figure 5b-d**). To quantify the time window over which the spiking activity of individual neurons is related to past, present, and future self-motion states, we compared the width parameters of model fits to decoding latency curves between the two behavioral tasks (**Figure 5h-i**). As predicted from qualitative observations, the temporal window for accurate linear speed decoding was significantly longer in duration during pursuit when compared to free explore (**Figure 5h**; Wilcoxon sign rank test, n = 148, median Pursuit – FE = 280ms, IQR = 0.00ms-523ms, z = −6.97, p = 3.13×10^−12^). The decoding window for angular speed was similarly extended during pursuit despite overall greater decoding accuracy during free explore (**Figure 5i**; Wilcoxon sign rank test, n = 86, median Pursuit – FE = 228ms, IQR = 85 – 427ms, z = −6.69, p = 2.31×10^−11^).

The duration of the decoding window may arise as an artifact of the increased continuity of behavior during pursuit (**sFigure 3c-d**). Numerous aspects of the data suggest this is not the case. First, this continuity was only statistically greater in pursuit for linear speed (**sFigure 3c-d)**, yet the duration of decoding was temporally extended for both angular and linear speed during target chasing. If the extended window was epiphenomenal to behavior, we would not expect to also observe it for angular speed decoding. Further, behavioral similarity over extended temporal windows was primarily driven by the presence of trial-like structure during pursuit which resulted in the animal spending more time immobile than during foraging (**sFigure 3**). To explore this possibility, we again analyzed spike train shifted decoding after excluding time frames in which the animal was not moving (speed < 5ms). Longer windows of self-motion decoding during target pursuit were again observed when differences in immobility were accounted for (Wilcoxon sign rank test, linear speed, n = 152, median Pursuit – FE = 149ms, IQR = −40-402ms, z = −5.81, p = 6.21×10^−9^; angular speed, n = 86, median Pursuit – FE = 206ms, IQR = 80 – 366ms, z = −6.75, p = 1.48×10^−11^).

Collectively, these results indicate that parietal cortex neurons tend to track the previous self-motion state of the animal irrespective of navigational context, yet exhibit self-motion related activation over elongated temporal durations during increased navigational demands (i.e. target chasing). This latter, and perhaps most striking result from self-motion decoding analyses, potentially provides a coding mechanism by which individual neurons can integrate past, current, and future movements of the animal to facilitate behavior-related prediction and path integration.

### Parietal cortex tracks visual target position in egocentric coordinates

Having demonstrated that tuning of PPC neurons to self-motion is enhanced and spans a longer integration window during pursuit, we next examined whether and how the pursuit target itself is integrated into PPC firing patterns. Pursuit requires efficient coordination of movement behavior in response to the position of the visual target relative to the animal. We hypothesized that neurons in PPC would be responsive to the bearing and/or distance of the target in egocentric coordinates. However, accurate detection of neurons that track the position of the visual target relative to the animal is complicated by the fact that pursuit behavior requires simultaneous covariation of spatial and self-motion variables that are known to modulate activation in some cortical neurons. Thus, sensitivity to visual target position could in actuality be explained by coupled relationships between target location and the aforementioned covariates. For instance, a neuron with sensitivity to the target occupying space to the animal’s left could potentially be explained by firing related to counterclockwise orienting behaviors that are more likely to occur in this scenario.

Accordingly, we implemented a generalized linear model (GLM) to examine the relative influence of spatial (position and head direction), self-motion (angular and linear speed), and target position (egocentric bearing and distance to the visual target) on the probability of spiking (**Figure 6a**). We also included predictors for egocentric boundary vector (EBC) related tuning, as neurons in PPC have recently been shown to exhibit activation in response to the distance and angle of nearby boundaries (Gofman et al., 2019; Alexander et al., 2020). For each neuron we constructed a model using all spatial, self-motion, EBC, and egocentric target position predictors and then examined the influence of each predictor by removing it from the model and assessing whether it significantly impacted fit by comparing the difference in log-likelihood of the reduced model (relative to the full model) to a chi-square distribution.

The GLM framework revealed that many PPC neurons were significantly modulated by combinations of behavioral and spatial predictors (**Figure 6b**). Consistent with the analyses above, large proportions of cells were sensitive to linear and angular speed. GLM analysis also demonstrated that large percentages of PPC neurons possessed sensitivity to the position of boundaries in egocentric coordinates relative to the animal, replicating the presence of egocentric boundary vector cells in these areas (**sFigure7a-b**). Many neurons were sensitive to head direction and allocentric position predictors in PPC. Additional experiments are needed to assess whether stable spatial correlates in this region are truly referenced to the allocentric reference frame (i.e. distal cue rotations), and it is important to consider that the proportion of cells sensitive to the these variables may be inflated by the presence of: 1) egocentric boundary cells which have spatially anchored firing as a consequence of activation relating to egocentric distance and orientation to boundaries and 2) characteristic trajectories in pursuit which couple self-motion behavior and reliable rat-to-target relationships with specific locations in space (note that proportions of HD and position sensitive neurons are increased during target chasing relative to free exploration).

33% of PPC neurons (n = 100/302) had significant decrements to model fit when the predictor for egocentric visual target position was dropped from the model (**Figure 6b,** ‘VT’). To visualize the responses of target sensitive neurons we next computed rat-to-target egocentric ratemaps. For each spike and position frame from each neuron we calculated the bearing and distance to the visual target relative to the animal in egocentric coordinates (centered on the rat’s current head direction) akin to analyses of behavior in **Figure 1**. We then generated rat-to-target position ratemaps for each neuron by normalizing the binned spike count for each location of the target (20° angular bins, 5 cm distance bins) by the amount of time the target occupied that bearing and distance (**Figure 6c**). The maximum target distance included in the analysis was restricted to 40cm which was qualitatively assessed to be where target occupancy dropped off precipitously (**Figure 6c, 1d-e**).

PPC neurons identified as having significant tuning to the target typically had robust target position receptive fields that fell into one of two forms: 1) restricted receptive fields possessing both a bearing and distance component (**Figure 6c, Figure 6d**, bottom rows) or 2) broader receptive fields primarily sensitive to egocentric bearing to target (**Figure 6d, top row**). Preferred bearings were distributed bilaterally with an average tuning width of approximately 70° (**Figure 6e**). The distribution of preferred distances fell within two modes ranging from animal proximal (<20cm) to more distal (>30cm) potentially reflecting the two forms of target-to-rat receptive fields. Several PPC neurons had preferred bearings that were behind the animal. However, the distribution of preferred bearings was skewed in front of the animal when the population of neurons with significant target sensitivity was restricted to those with reliable bearing tuning across non-overlapping epochs within the target chasing session (**Figure 6e**, dark blue histograms). Accordingly, preferred bearings behind the animal likely reflect the influence of other covariates on rat-to-target ratemaps (e.g. actions, the position of boundaries, etc.).

Indeed, many neurons with sensitivity to the position of the visual target were conjunctively modulated by other predictors including the egocentric position of arena boundaries (**Figure 6b**). Approximately half of neurons with egocentric boundary correlates were also modulated by the egocentric position of the visual target suggesting that neurons with this coding property can map and integrate the egocentric position of multiple environmental features simultaneously (**sFigure7c**). Further, nearly half of neurons with self-motion sensitivity also responded to the position of the target relative to the animal (**Figure 6f**).

Finally, we suspected that multiplicative gain modulation of self-motion related activation could emerge in neurons that simultaneously tracked the visual target. In support of this hypothesis, linear speed sensitive neurons with target sensitivity had significantly greater differences in within-receptive field amplitude (i.e. multiplicative gain) and mean rate (i.e. additive gain) between pursuit and free explore than the subset of neurons that did not possess responsivity to the target (**Figure 6g**, Linear speed amplitude difference; VT, n = 65, median = 1.02Hz, IQR = 0.50 – 1.85Hz; no VT, n = 88, median = 0.57Hz, IQR = 0.27 – 0.93Hz; Kolmogorov-Smirnov (KS) test, D = 0.32, p = 8.46×10^− 4^; Linear speed mean difference; VT, median = 0.86Hz, IQR = 0.41 – 1.16Hz; no VT, n = 88, median = 0.49Hz, IQR = 0.16 – 0.98Hz; KS test, D = 0.27, p = 0.008). No significant differences were observed between the populations of angular velocity sensitive neurons with or without visual target sensitivity (**Figure 6g**, Angular speed amplitude difference, D = 0.14, p – 0.81; mean difference, D = 0.19, p = 0.44). These results indicated that gain modulation of linear speed tuning likely arose from the multiplexing of movement commands with information relating to the target position in egocentric coordinates.

## Discussion

We developed an ecologically inspired paradigm wherein rats were trained to pursue moving visual targets in a manner akin to hunting or social chase behaviors observed in non-laboratory settings (Calhoun, 1963). From a behavioral standpoint, we established that 1) rats can be trained to chase purely visual stimuli that are associated with reward and, 2) rats exhibit predictive processing which manifested as shortcutting routes that intercepted the target when it traversed familiar trajectories. We also demonstrated that self-motion tuning in PPC is impacted by navigational context. Specifically, the relationship between spiking activity and self-motion tuning was more nonlinear and reliable during target pursuit than free exploration. These changes to tuning functions resulted in greater numbers of significantly self-motion tuned neurons in PPC during pursuit. Further, PPC self-motion tuning was subject to both additive and multiplicative gain modulation which enhanced decodability of self-motion related to the pursuit behavior. Finally, we showed that individual PPC neurons tracked both self-motion and the egocentric position of the visual target, and that neurons with this property exhibited greater gain modulation than those that did not. Collectively, these results indicate that PPC adaptively codes for multiple task-relevant idiothetic and external variables over temporal integration windows related to the statistics of goal-directed behavior.

### Predictive processing, shortcuts, and memory

Pursuit behaviors demand continuous tracking of a target and mirroring of its trajectories. We observed both behavioral components during performance of pseudorandom trials in the present study (**Figure 1, sFigure 1**). If the trajectory of the target can be predicted based on previous experience, pursuit behaviors may also include attempts at target interception. Such behavior would require short-term memory processes involving motion integration and prediction. Indeed, locomotor behavior on probe trials using a ‘characteristic’ path having a stable shape and start and end points evidenced seconds-long integration of trajectories. This manifested as predictive ‘shortcutting’ behavior yielding interception of the target’s path and, neurophysiologically, as retrospectively lagged spike rate correlates to linear and angular velocity of self-motion (discussed further below).

Several lines of evidence suggest rats cannot infer novel routes in labyrnthian mazes and instead must learn to execute shortcut trajectories (Tolman, 1948; Grieves and Dudchenko, 2013). All data in the present study were collected after rats had reached asymptotic performance on the target chasing task and thus, we cannot examine the emergence of shortcutting behavior over experience.

That said, shortcutting routes were observed across recording sessions spanning weeks to months which indicates that the shape of characteristic trajectories were learned and stored in memory. Such learning occurred despite the fact that the full trajectories themselves could not be viewed in full at any given moment. Longer term memory for specific routes and their spatial relationships to environmental boundaries may be relevant in many navigational contexts including pursuit. For example, following behavior may be motivated by the need to learn a specific route between two locations and/or the need to update current location relative to a start, or ‘home’, location. Predictive behaviors in our data demonstrate that rats can learn route shapes and quickly recognize known trajectories of the target as a simple function of experience. The emergence of inferential behavior in our task establish a behavioral framework by which systems dynamics supporting predictive processing in freely moving rodents can be further examined.

### Adaptive PPC self-motion tuning and gain modulation

While several recent studies have examined sub-cortical mechanisms that appear to drive pursuit behaviors in rodents, we examined the PPC, which in navigating rodents exhibits conjunctive encoding of egocentric goal location, locomotor actions such as turning, and progress through a route (Chen et al., 1994; Whitlock et al., 2012; Wilber et al., 2014, 2017; Nitz, 2006, 2012). Consistent with previous reports, we observed that a majority of PPC neurons were tuned to linear and/or angular speed during both pursuit and foraging navigation, but that this tuning was dynamically modulated in several ways as a function of task (Whitlock et al., 2012). The profiles of movement related tuning took many forms including nonlinear functions as observed in structures such as the medial entorhinal cortex (MEC, (Hinman et al., 2016). Tuning functions became more nonlinear and reliable during pursuit which yielded greater proportions of neurons sensitive to self-motion when compared to free exploration (**Figure 2**).

PPC neurons also exhibited increased mean firing rate relationships to linear speed during chase behavior, even when controlling for self-motion differences between each locomotor regime (**Figure 2, sFigure 3–4**). This presented as additive gain modulation across the range of velocities examined. However, multiplicative gain modulation was also observed. As tuning curves for most neurons were nonlinear, we fit each curve with Gaussian-modified linear models and showed that multiplicative firing rate increases were primarily restricted to tuning curve peaks during pursuit (**Figure 3, sFigure 5**). This latter finding parallels multiplicative gain modulation observed in sensory cortices wherein multimodal contextual information increases the magnitude of response for an individual neuron without altering its base receptive field properties (e.g. an orientation selective visual cortex neuron will increase firing at its preferred orientation when the animal is mobile versus immobile; Ferguson and Cardin, 2020; Niell and Stryker, 2010; Salinas and Thier, 2000). We believe our data provide the first evidence of such gain modulation in relation to the coding for idiothetic variables in freely moving rats.

Beyond gain modulation, adaptation of self-motion tuning curves as a function of navigational context was seen in at least two other ways that were related to differences in movement between foraging and pursuit. During pursuit, movement of the visual target was smooth with few abrupt changes to angular velocity. To pursue efficiently, rats primarily needed to prioritize straightforward motion and match their linear speed with the target (**sFigure 3**). In keeping with these changes in the distributions of self-motion, shifts in the velocities associated with peak firing moved systematically toward the higher values observed more frequently during pursuit. This was observed even though we matched sampling of self-motion values across the pursuit and foraging conditions (**sFigure 3**). In addition to shifts in tuning curves to higher linear speeds, decoding of self-motion was enhanced for linear speed but not angular speed in pursuit, again matching shifts in behavior required to perform the task. Thus, adaptations in PPC self-motion tuning followed variation in how the animal moved within the environment between navigation tasks.

### Self-motion integration in PPC

Evidence for self-motion integration, as opposed to moment-to-moment encoding of self-motion was established by shifting spike trains relative to behavioral data and determining the accuracy of self-motion prediction (**Figure 5**). The distribution of optimal latencies between instantaneous firing rate and self-motion clearly favored retrospective tuning counter to reports of anticipatory responses in PPC and other regions (Moore et al., 2017; van Wijngaarden et al., 2020; Whitlock et al., 2012). These preferred latencies were stable between random foraging and pursuit sessions, suggesting that PPC neurons are strongly influenced by the history of self-motion, irrespective of behavioral task. This finding extends previous work showing that PPC is critical for decision making based on prior trial outcomes to the domain of navigation (Morcos and Harvey, 2016; Hwang et al., 2017; Akrami et al., 2018). Further, retrospective coding of self-motion could support memory processes critical for the internalization of characteristic routes required for the execution of predictive shortcuts.

Across neurons, self-motion prediction accuracy exceeded chance for temporally lagged relationships between neural activity and behavior on the order of seconds extending both behind and ahead in time relative to ongoing behavior. Temporally extended windows of accurate linear speed decoding have also recently been reported in the medial entorhinal cortex, suggesting that movement tracking at behaviorally-relevant timescales is a general coding property of cortex (Dannenberg et al., 2019). PPC neuronal activity exhibited these temporally extended relationships to self-motion in both free exploration and pursuit, and since this same population of neurons exhibits tuning that spans all combinations of linear and angular velocity values, it follows that a stable population of PPC neurons together reflect the past, present, and future trajectory of the animal.

Such time-lagged tuning of locomotor states perhaps explains why PPC ensembles produce unique firing patterns for all positions along a complex path even when specific actions such as left/right turns and directions of travel repeat multiple times across the same route (Nitz, 2006, 2012). Under such conditions, route positions sharing the same momentary linear and angular velocity combination will nevertheless differ in the linear and angular velocity sequences that precede them. Time-lagged tuning to self-motion parameters could also underlie the development of distance and sub-space trajectory encoding during track-running behaviors in retrosplenial cortex which receives dense innervation from PPC (Alexander and Nitz, 2017). This extended integration window could associate head directions, allocentric and within-route locations, and egocentric views during navigation to produce conjunctive spatial representations observed in these association regions that are thought to mediate egocentric–allocentric reference frame transformations (Cohen and Andersen, 2002; Byrne et al., 2007; Nitz, 2012; Smith et al., 2012; Wilber et al., 2014; Alexander and Nitz, 2015; Vedder et al., 2016; Krumin et al., 2018; Minderer et al., 2019).

Critically, integration of self-motion was adaptive to the locomotor regime of the animal. While the optimal temporal offsets at which individual neurons most precisely encoded self-motion was stable across foraging and pursuit, the time scale over which effective tuning was observed was adaptive. By examining autocorrelations for linear and angular velocity traces across time, we were able to quantify the extent of time over which linear and angular velocities persisted. Both were longer under conditions of pursuit as compared to foraging. Under pursuit, animals exhibited longer bouts of forward running with smaller variation in angular velocity. In parallel, the scale of time over which any given neuron’s activity was effectively tuned to linear and angular velocity sequences was extended under pursuit. In this way, the tuning properties of PPC neurons were found to be adaptive to the distribution of locomotor variables in different locomotor regimes. We speculate that such adaptation allows the system to encode locomotor sequences and their associated trajectories over a range of distances per unit time (Andersen and Cui, 2009). Similar observations of extended integration timescales have been reported in PPC during evidence accumulation in head-fixed virtual navigation tasks, and we replicate and extend these observations in two ways (Morcos and Harvey, 2016; Runyan et al., 2017). First, we show that the longer integration windows in PPC are not restricted to behaviors requiring explicit evidence accumulation and decision making and instead emerge during natural foraging navigation. Second, we show that the timescale is adaptive to the statistics of behavior which, in our case, is reflected in the increased decoding timescale during pursuit to match alterations to movement between conditions.

### Mechanisms of PPC adaptation and efferent targets

Together, the results indicate that pursuit, as compared to random foraging, is associated with higher mean firing rates (i.e. additive gain) in PPC, higher peak firing rates (i.e. multiplicative gain), enhanced reliability in firing rate as a function of linear and angular velocity, greater numbers of self-motion sensitive neurons, systematic shifts in the velocities associated with peak firing, and increased accuracy of self-motion decoding over longer time windows of self-motion integration. Context-invariance of linear speed coding has been reported in the MEC so these observations may be unique to PPC or other association areas receiving multisensory input (Kropff et al., 2015). Future work will be required to address what specific network dynamics facilitate the adaptations to PPC idiothetic processing observed in response to navigational context.

The most obvious sensory difference between pursuit and free foraging is the presence of the visual target. Our neuronal recordings were conducted in medial PPC (mPPC/V2m) where there is less direct innervation from primary visual cortex or thalamic visual areas (Reep et al., 1994; Nitz, 2009; Wilber et al., 2015; Olsen and Witter, 2016; Olsen et al., 2017). However, it is important to note that analogous anatomical regions in mice are considered part of the higher visual cortex (Glickfeld and Olsen, 2017; Hovde et al., 2019; Gilissen et al., 2020) and that mPPC receives dense afferents from dysgranular RSC (dRSC) where visually-evoked responsivity has been reported (Fischer et al., 2020; Mao et al., 2020; Powell et al., 2020; Zhuang et al., 2017). Taken together, it seemed highly probable that PPC activity would be influenced by visual information and consistent with this hypothesis, we report that a subset of PPC neurons were sensitive to the egocentric position of the visual target (**Figure 6**). Accordingly, the observed changes to firing rate gain could result purely from increased activity relating to the presence of a visual stimulus. The observed gain modulation could in turn alter reliability of self-motion tuning, signal-to-noise ratios, and consequently, decodability of linear speed. As the stimulus is present for the entirety of a pursuit trial, persistent visually-driven excitation of PPC ensembles could underlie the temporally extended windows of self-motion decoding observed during chase. Additional studies should test the role of visual processing streams on the observed adaptations to self-motion processing in PPC during pursuit behavior.

Beyond the influence of purely visually-evoked excitation, the target dictates behavior and undoubtedly engages arousal and attentional mechanisms known to modulate cortical activation and facilitate gain modulation (Ferguson and Cardin, 2020; Reynolds et al., 2000; Vinck et al., 2015). Visual attention and overall arousal state modulate activation of PPC neurons in multiple species (Bucci et al., 1998; Colby and Goldberg, 1999; Tingley et al., 2014; Stitt et al., 2018). Indeed, PPC neurons with simultaneous sensitivity to target position and linear speed exhibited the strongest additive and multiplicative gain modulation. This enhancement may be facilitated by activation of the basal forebrain (BF) which, among other neuromodulatory projections, forms cholinergic afferents into cortex thought to be critical for gain modulation relating to sensory processing (Disney et al., 2007; Goard and Dan, 2009; Minces et al., 2017; Záborszky et al., 2018). BF connectivity forms reciprocal loops with neocortical regions, including the PPC (Bigl et al., 1982; Gritti et al., 1997; Zaborszky et al., 2015). Accordingly, it will be important to investigate whether PPC sensitivity to the position of the visual target signals to the BF to enhance the response gain. Conversely, BF activation relating to increased arousal or attention to the target could alter the input threshold required for PPC spiking and consequently enable neurons to form egocentric receptive fields for the position of the visual target. Regardless, neuromodulation likely underlies gain modulation of self-motion processing observed during pursuit sessions in the present work. Other cellular mechanisms of gain modulation are also likely involved including alteration to GABAergic inhibitory circuits and modulation of corticothalamic loops (Fu et al., 2014; Ferguson and Cardin, 2020).

We conclude that changes to PPC dynamics during pursuit illustrate systematic and flexible processing recruitment in the service of behavioral demands associated with a specific navigational context. This adaptation alters the temporal scale by which PPC processes information, perhaps to specifically organize movement sequences as a function of task. These task-specific changes to self-motion processing could influence motor coding observed in the M2 region of frontal cortex where neurons are sensitive to the context of actions (Reep et al., 1994; Nitz, 2012; Olson et al., 2020). Pursuit related changes to PPC self-motion computations appear ideal for coordinating movement and trajectories at multiple timescales in response to the position of visual targets and may complement the pursuit related dynamics in the striatum, superior colliculus, and hypothalamus by enabling predictive behaviors (Schiller and Stryker, 1972; Cooper et al., 1998; Hoy et al., 2016, 2019; Shang et al., 2019; Zhao et al., 2019; Kim et al., 2019).

## Methods

### Subjects

Male Long-Evans rats (n = 6) served as behavioral subjects and were housed individually and kept on a 12-h light/dark cycle. Animals were habituated to the colony room and handled daily for a period of 1-2 weeks prior to training on the visual target chasing task. Rats were food restricted to approximately 85-90% of their free-fed weight. Water was available continuously. All experimental protocols adhered to AALAC guidelines and were approved by IACUC and the UCSD Animal Care Program.

### Target chasing behavior

Animals were trained to pursue a moving light stimulus in a 4ft diameter circular arena for reward. The visual stimulus was a 1.25 cm dot from a bright green laser pointer controlled by one of two experimenters in the experimental room. The arena was placed on a table 3ft above the ground. Boundaries of the arena were 2.5cm in height and fixed distal cues were outside of the arena across all sessions. Prior to the initiation of training all rats were handled extensively.

Acquisition of visual target pursuit behavior required behavioral shaping over approximately 1 – 2 months of training. Rats were initially habituated to the arena for 1 week by randomly placing cereal bits around the arena and allowing the animal to free forage for 20 minutes per day. After arena habituation the visual stimulus was introduced. During this stage, 5-10 cereal pieces would be present on the arena at any given time and the experimenter would direct the visual stimulus to hover on top of rewards that the animal was about to ingest. After the animal acquired a reward that the light stimulus was positioned on, the stimulus would shut off in order to create an association between the visual target and reward. After approximately 1 week of this process the next phase of shaping began. In the third stage, a single cereal piece was tossed to a random position in the arena at a time. The visual stimulus hovered over the reward and shut off when the animal retrieved it. This step repeated continually for a 20-30-minute session for approximately 1-2 weeks or until the animal was readily running to the visual target/reward position. In the final shaping phase, the onset of the stimulus would occur in the absence of a reward in the arena. Because the visual target was associated with reward, the animal would approach it. Prior to the rat reaching the stimulus, the experimenter would move the visual target smoothly away from the animal and in most cases the rat would engage in pursuit. If the rat intercepted the stimulus it would disappear, and a reward would be tossed into a random location within the arena. The visual target would then reappear hovering over the reward location, and again disappear after the animal retrieved the reward. Following reward consumption, the next trial would begin by activating the visual stimulus in a pseudorandom location within the arena and the aforementioned process would repeat. Over approximately a 1-month period (duration was dependent on the behavior of individual rats), the length of pursuit could be extended without causing the animal to lose interest or become frustrated.

After shaping, rats would chase the visual target for approximately 25 minutes per day during concurrent in vivo electrophysiological recordings. The visual target primarily moved in pseudorandom trajectories (RTs) but characteristic trajectories (CTs) of the target were also instantiated after animals were frequently chasing the visual target. For CTs, the light target would execute a stereotypical path that began and ended at the same approximate allocentric locations within the arena and recording room. CTs occurred less frequently than RTs and were randomly interspersed throughout the target pursuit block. Two CTs were utilized across animals. CT1 moved from north to south and required the animal to execute a clockwise U-Turn while CT2 moved from east to west and required a counterclockwise U-Turn (**sFigure 1**). Target pursuit blocks could occur before or after free foraging sessions conducted in the same arena (approximately 10-20 minutes). In a subset of recordings, the pursuit session was conducted between two free foraging sessions of approximately 10-15 minutes each.

### Surgery

After readily engaging in 20+ pursuits of the visual target in a session, rats were surgically implanted with tetrode arrays (twisted sets of four 12 micrometer tungsten wires or 17 micrometer platinum-iridium wires) fitted to custom-fabricated microdrives that allowed movement in 40μm increments. Each microdrive contained 4 tetrodes. Rats were implanted with 3 total microdrives positioned in either posterior parietal cortex (PPC) or retrosplenial cortex (RSC, data not shown). Rats were anesthetized with isoflurane and positioned in a stereotaxic device (Kopf Instruments). Following craniotomy and resection of dura mater overlying cortical regions, microdrives were implanted relative to bregma (PPC, A/P −3.8 mm, M/L ± 2.2 mm, D/V −0.5 mm; RSC, A/P −5.8 mm, M/L ± 0.7-1.2 mm, D/V −0.5 mm, 10-12° medial/lateral angle). 1 of the 6 rats trained on the behavior did not have a PPC implant and is therefore included in behavioral analyses (**Figure 1**) but not included in analyses of neural data (remaining figures).

### Recordings

Each microdrive had one or two electrical interface boards (EIB-16, Neuralynx) individually connected to amplifying headstages (20X, Triangle Biosystems). Signals were initially amplified and filtered (50x, 150Hz) on the way to an acquisition computer running Plexon SortClient software. Here the signal was digitized at 40kHz, filtered at 0.45-9 kHz and amplified 1-15X (to reach a total of 1,000-15,000X). Electrodes were moved ventrally (40μm) between recordings to maximize the number of distinct units collected for each animal. Single-units were manually identified using Plexon OfflineSorter software. Primary waveform parameters utilized were peak height, peak-valley, energy, and principal components.

Animal position was tracked using a camera set 10ft above the recording room floor. Plexon’s CinePlex Studio software was utilized to separately detect blue and red LED lights. Lights sat approximately 4.5 cm apart and were positioned perpendicular to the length of the animal’s head. During the target chasing paradigm, the green visual stimulus was simultaneously tracked using the same software.

### Histology

Animals were perfused with 4% paraformaldehyde under deep anesthesia. Brains were removed and sliced into 50μm sections and Nissl-stained to identify the trajectory and depth of electrode wires in PPC and RSC. Boundaries of each region were defined in accordance with our previous work as well as the Paxinos and Watson and Zilles atlases (Paxinos and Watson, 2006; Zilles, 2012). All tetrodes were determined to be within the bounds of mPPC. Documented micro-drive depth across recordings and final electrode depth observed in histology were compared and found to be compatible in all cases.

### Analyses

#### Identification of pursuits and shortcuts

Pseudorandom (RTs), characteristic (CTs), and shortcut trials were identified using a custom graphical user interface (GUI) designed in MATLAB. The GUI enabled fine resolution scoring of starts and ends of runs. Starts of runs were identified as timepoints wherein the visual target and rat began moving coherently. Ends of runs were identified as timepoints wherein the rat intercepted the visual target and the visual target disappeared (i.e. tracking of the target was lost). CTs were easily identifiable relative to RTs because of their consistent shape and position in the environment. The total amount of CTs identified using tracking data were cross-referenced with recording logs from each session. Shortcuts were also readily identified, as they were similar to CTs but the animal intercepted the target at an earlier point and the target disappeared. Median egocentric bearing and distance (described below) across all position samples from all RTs, CTs, and shortcuts were compared and verified identification of the different trial types.

#### Latency between position of rat and visual target

To assess the temporal proximity of rat pursuit of the visual target between pseudorandom trajectories (RTs) and characteristic trajectories (CTs) the x- and y-position of the animal was concatenated and shifted in time relative to the position of the visual target in 50ms increments and similarity was assessed via Spearman’s correlation (ρ) at each lag. Latency was defined as the shift in rat position that yielded maximal correlation. This often required shifting backwards in time to match the rat position to the prior position of the target (e.g. the animal is now at the same position that the visual target was 150ms ago).

#### Autocorrelations of movement variables

To assess differences in movement statistics between free exploration (FE) and pursuit (P) we calculated autocorrelations (‘xcorr’ in MATLAB) of linear and angular speed (**sFigure 3c**). To determine the temporal continuity of movement behavior we found the temporal lag wherein the autocorrelation dropped below 0.25 for each recording session. Rats exhibited more similar linear speed for extended temporal windows during P when compared to FE. Continuity of angular speed was more similar between FE and P, but FE had significantly greater temporal lags (**sFigure 3d**).

#### Analysis of the egocentric position of the visual target position relative to the animal

For each position frame of each recording session, the instantaneous distance between the rat and the visual target was calculated as the Euclidean distance in centimeters (cms) between the average x- and y- positions of tracking lights on the animal and the tracked position of the target. The instantaneous egocentric bearing to the visual target was calculated as the difference between the rat’s heading direction and the inverse tangent of the difference between x- and y-positions of the visual target and the rat.

To examine the distribution of target occupancy relative to the animal, occupancy maps were generated by binning egocentric distance in 2.5cm increments (up to 65cms) and egocentric bearing in 5° angular bins. Differences in egocentric target occupancy between RTs and CTs were assessed by generating target occupancy maps separately for each trial type. We summated across distance and bearing bins to create marginalized occupancy vectors for egocentric bearing and distance separately, then fit these with Gaussian-modified linear models (GML) using the built-in *‘fit’* function in MATLAB. As the GML model is utilized in multiple analyses throughout the paper we define it now. GMLs took the form:

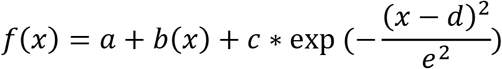

Where *a* and *b* are the intercept and slope of the linear fit and *c, d,* and *e* are the amplitude, center, and standard deviation of the additive Gaussian function. To quantify differences in target distance, bearing and bearing deviation, we compared the center and standard deviations of model fits between RTs and CTs.

#### Construction of self-motion tuning curves and normalization of behavior free foraging and pursuits

Behavior-matched angular and linear speed tuning curves were constructed for all neurons. Within a recording session, we discretized linear speed into 29 bins spanning 0 to 45 cm/s (1.6 cm/s bins) for free exploration (FE) and target pursuit (P) separately. Angular speed was similarly binned (34 bins ranging from −180° to +180° in 10.6° increments). For each self-motion variable, the minimum occupation across all speed bins across both FE and P epochs was identified. We next computed 1000 tuning curves for each individual neuron for P and FE(s), each time sub-sampling from all possible spike train indices (within P or FE) at a given speed bin to match this minimum occupation time minus an additional 1 second. All tuning curves depicted in the manuscript are the mean (± s.e.) of these 1000 sub-sampled tuning curves. Using this method all self-motion tuning curves were behaviorally matched between FE and P as well as across speed bins. We repeated this process for each neuron after randomly shifting the spike train relative to self-motion behavior (within FE and P blocks) 1000 times to generate random (null) self-motion tuning curves. Reliability of tuning curves was assessed within a behavioral epoch by repeating this process but for non-overlapping odd and even minutes within a session, then assessing their similarity using Spearman’s correlation (ρ).

#### Fitting self-motion tuning curves, model comparisons, and identification of significant tuning

Each tuning curve for each neuron was fit with a uniform function, linear fit, and the GML model described above. All randomized tuning curves for each neuron were also fit with these models. F-tests were run to compare the residuals of all combinations of model fits. Any neuron with a true p-value for either uniform versus linear or uniform versus GML fits that was less than the 1^st^ percentile of the same tests conducted on the neuron’s corresponding null self-motion tuning curves was considered to have significantly nonuniform tuning to self-motion. The final subset of neurons identified to have significant self-motion sensitivity passed this metric for at least one session (i.e. FE1, P, or FE2) and had a reliability score (Spearman’s ρ, as described above) that was greater than the 99^th^ percentile of the distributions of reliability scores calculated from each neuron’s null self-motion tuning curves. Significant nonlinearity of tuning curves was attributed to neurons with p-values from F-tests of linear versus GML fits that were less than the 1^st^ percentile of the distributions of p-values computed from similar tests conducted on model fits to null self-motion tuning curves. All percentile tests were done within neuron and behavioral task (i.e. from the distribution of p-values on model fits to all 1000 of the null self-motion tuning curves calculated from randomly shifting that neuron’s spike train within pursuit or free exploration).

#### Assessment of gain modulation

Additive gain in self-motion tuned neurons for FE or P was determined by peak normalizing self-motion tuning curves for each epoch by the maximum firing rate value across tuning curves taken from all behavioral sessions, then taking the mean difference between these normalized tuning curves between P and FE. Neurons with mean differences less than zero prefer FE (P<FE) while neurons mean differences above zero prefer P (P>FE). To assess multiplicative gain, we examined differences in the amplitude parameter of GML model fits to tuning curves between P and FE. Systematic modulation of both additive and multiplicative gain was assessed by comparing differences in mean rate or peak amplitude between all combinations of P, FE1, and FE2 for the subset of self-motion tuned neurons recorded in two FE conditions.

#### Decoding of self-motion

Decoding of self-motion at the level of single neurons was conducted using a maximum correlation coefficient classifier taken from the Neural Decoding Toolbox (Meyers, 2013); http://www.readout.info/). Spike times for a given neuron were matched to behavioral position samples to generate a spike train with the same temporal resolution as the position tracking system (60Hz). Spike trains were smoothed with a Gaussian filter with a 200ms standard deviation and partitioned into 50% training and 50% test datasets. The classifier was trained on the instantaneous firing rate vector with self-motion (either linear or angular speed) as the response variable. Self-motion was binned in the same manner as for analysis of tuning curves described above. The training data was used to generate a template by learning the mean firing rate associated with each class (i.e. speed). Predictions on test data are chosen by finding the class with the smallest square difference between the template and a given test sample (i.e. a single firing rate bin in the spike train). Decoding was performed with 10-fold cross validation using different 50/50 randomly selected splits of train and test data and accuracy was assessed via Spearman’s correlation between predicted and true self-motion. All correlations reported are the average across cross validations.

Decoding of self-motion using simultaneously recorded PPC ensembles was conducted with a naïve Bayes classifier again using the Neural Decoding Toolbox. Ensembles ranged in size from 5 to 12 PPC neurons and the classifier was trained on the same spike trains generated for single neuron prediction and self-motion response variables. Training data was used to calculate the discretized mean firing rates for each neuron for each class (i.e. speed bin) and the log likelihood function is calculated using these rates as lambda parameters to define Poisson distributions for each neuron. For a single test point, the probability of observing the combination of mean firing rates across the ensemble is calculated for each class and multiplied across all neurons to give a likelihood of being at each speed. The speed with the highest likelihood is the predicted self-motion state. Data was partitioned in the same manner as outlined above with 10-fold cross validation and prediction accuracy was again assessed using Spearman’s correlations and averaged across folds.

#### Self-motion decoding latency analysis

To quantify the temporal relationship between neural activation and self-motion decoding, we additionally ran the maximum correlation coefficient classifier analysis for single neurons after *shifting the spike train* relative to behavior incrementally. Specifically, the spike train was shifted from −2s to +2s in 100ms increments relative to a fixed behavioral response variable (e.g. linear or angular speed). The shifted variable is key to interpretation as the temporal relationships relative the reference vector dictate whether lagged relationships are prospective (i.e. anticipatory) or retrospective (i.e. history-dependent). Because we shifted the spike train, backwards shifts relative to behavior mean spiking occurred *after* the response variable (i.e. speed), while forward shifts relative to behavior mean spiking occurred *before* the response variable. Increased decoding accuracy for the former indicates history-dependent relationships between neural activity and behavior while the latter evidences anticipatory relationships.

For each temporal lag iteration, the decoder was trained and tested as described above. Accuracy was again assessed by correlating the self-motion prediction associated with each spike train lag with the true self-motion. These values were stored in a 41-bin vector for each neuron which we refer to as the ‘Decoding accuracy latency curve.’ These curves were fit with the GML model and center and width parameters were extracted to assess the preferred latency and decoder width for each neuron, respectively. For all decoding analyses, including the instantaneous decoding described above, real decoding accuracy was compared to the 99^th^ percentiles of decoding accuracy taken from a null distribution for each neuron following circular shifts of the spike train relative to self-motion for durations greater than 2 seconds for 41 iterations (the number of spike train shifts in the latency analysis).

#### Generalized linear models (GLMs)

To test the influence of multiple behavioral and spatial variables on the activity of PPC neurons simultaneously we utilized a GLM framework (https://github.com/hasselmonians/pippin). The probability of spiking in a given behavioral frame (60Hz) is described by an inhomogeneous Poisson process, where the spiking probability in a given position sample is described by the continuous variable λ:

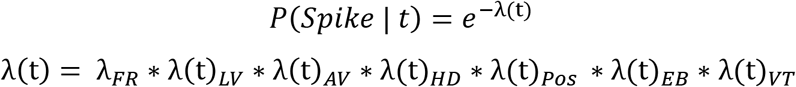

Where:

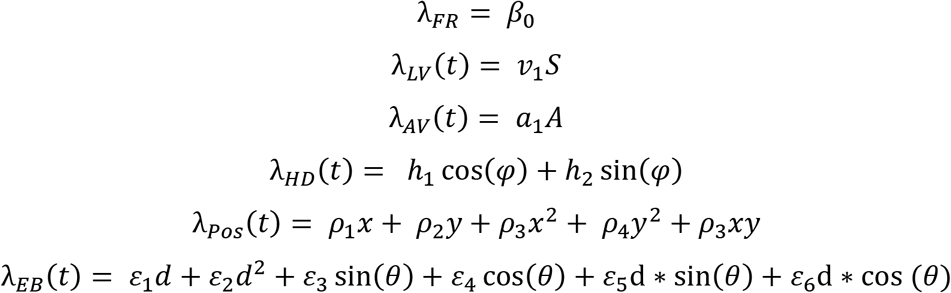

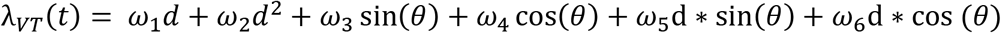

Where β_0_ defines the baseline firing rate of the neuron. All subscripted variables are fit coefficients weighting the other (time-varying) variables. S is the running speed of the animal and A is the angular displacement of the animal, as described above. φ is the head direction, and x and y are measurements of the animal’s position in the environment in centimeters. Finally, d is the animal’s distance from the center of the environment (EB) or the distance to the visual target (VT), and θ is the egocentric angle to the center of the environment (EB) or the egocentric angle to the visual target (VT). VT predictors are not included in GLMs for FE as no target was present.

Coefficients were determined by fitting to maximize log-likelihood (MATLAB function ‘glmfit’) of the experimental spike train given the behavioral variables. Model selection was completed in a stepwise fashion in order to identify the simplest model. On each iteration, we added the predictor which increased model fit (on average of 5-folds) the most, and was statistically significant compared to a Chi-Square distribution (degrees of freedom equal to the number of coefficients set to zero, p<=0.001). We then dropped any predictor which no longer had a significant contribution (p>0.05). This process was repeated until the model converged. While theoretically the change in log-likelihood should follow a Chi-Square distribution, this is only the case when the spike train has been fit well (e.g. including all neuron-neuron coupling terms). We therefore also compared the change in log-likelihood to that from 1000 randomly shuffled spike trains, giving an empirical null-distribution and results were similar to the Chi-Square approach. Example spike trains for each model were generated by evaluating lambda for each behavioral time point (‘glmeval’ in MATLAB) and using this as the input to a random Poisson Generator (‘poissrnd’ in MATLAB).

#### Rat-to-target ratemaps and quantification

To visualize and characterize the receptive fields of neurons with significant tuning to the egocentric position of the target using the GLM we created rat-to-target ratemaps. Rat-to-target ratemaps were constructed by finding the egocentric distance and bearing to the target at the time of all spikes for a given neuron within the pursuit session (see section on egocentric target position above). These values were used to generate spike-target-occupancy maps which were normalized by the time the visual target spent in each egocentric distance by bearing bin to generate the rat-to-target ratemaps depicted in **Figure 6**. As laser occupancy drastically dropped at 40cms away from the animal, the radial axis on rat-to-target ratemaps was restricted to values below this distance. Rat-to-target ratemaps were smoothed with a Gaussian kernel with a 5cm x 20° standard deviation.

Preferred bearing, tuning width, and preferred distance of cells determined to have significant sensitivity to the egocentric position of the visual target using the generalized linear model were quantified in a similar fashion to the target occupancy maps, again using parameters of GML fits. We examined these characteristics for all neurons determined to possess sensitivity to the egocentric position of the target using the generalized linear model as well as the distributions for neurons that were significantly sensitive and possessed reliability in the strength (MRL) and direction of their visual target related tuning across non-overlapping halves of the pursuit session (blue distributions in **Figure 6e**).

#### Egocentric boundary ratemaps and 2D spatial ratemaps

Egocentric boundary ratemaps (EBRs) were constructed as previously described using previously published code (https://github.com/hasselmonians/EgocentricBoundaryCells; Hinman et al., 2019; Alexander et al., 2020). In brief, the egocentric bearing and distance to all boundaries are calculated for all behavioral position samples and spike times for a given neuron. These values are utilized to create boundary and spike-boundary occupancy maps which reflect the position of the boundaries (<62.5cms from the animal) relative to all animal position samples within a session and relative to all times in which a spike occurred for a given neuron. Egocentric bearing is anchored to the allocentric head direction of the animal at every behavioral sample such that the animal is facing upwards in the polar matrices containing this occupancy information. EBRs are constructed for each cell by normalizing the spike-boundary occupancy map by the amount of time each egocentric boundary distance by bearing bin was occupied as shown in **sFigure 7**. Ratemaps were smoothed by convolving EBRs with a 2.5cm x 3° Gaussian kernel.

2D spatial ratemaps were constructed in a similar manner as above but using the x- and y-position of the rat in centimeters relative to the external environment. Rat position was discretized into 3×3cm bins. For a given neuron, the rat position at the time of all spikes was determined to generate a spike occupancy map which was normalized by the total time each spatial position was occupied to generate a 2D spatial ratemap. Raw ratemaps were smoothed by convolving with a Gaussian kernel with a 6cm^2^ standard deviation.

## Supplemental Figure Legends

**Supplemental Figure 1.**
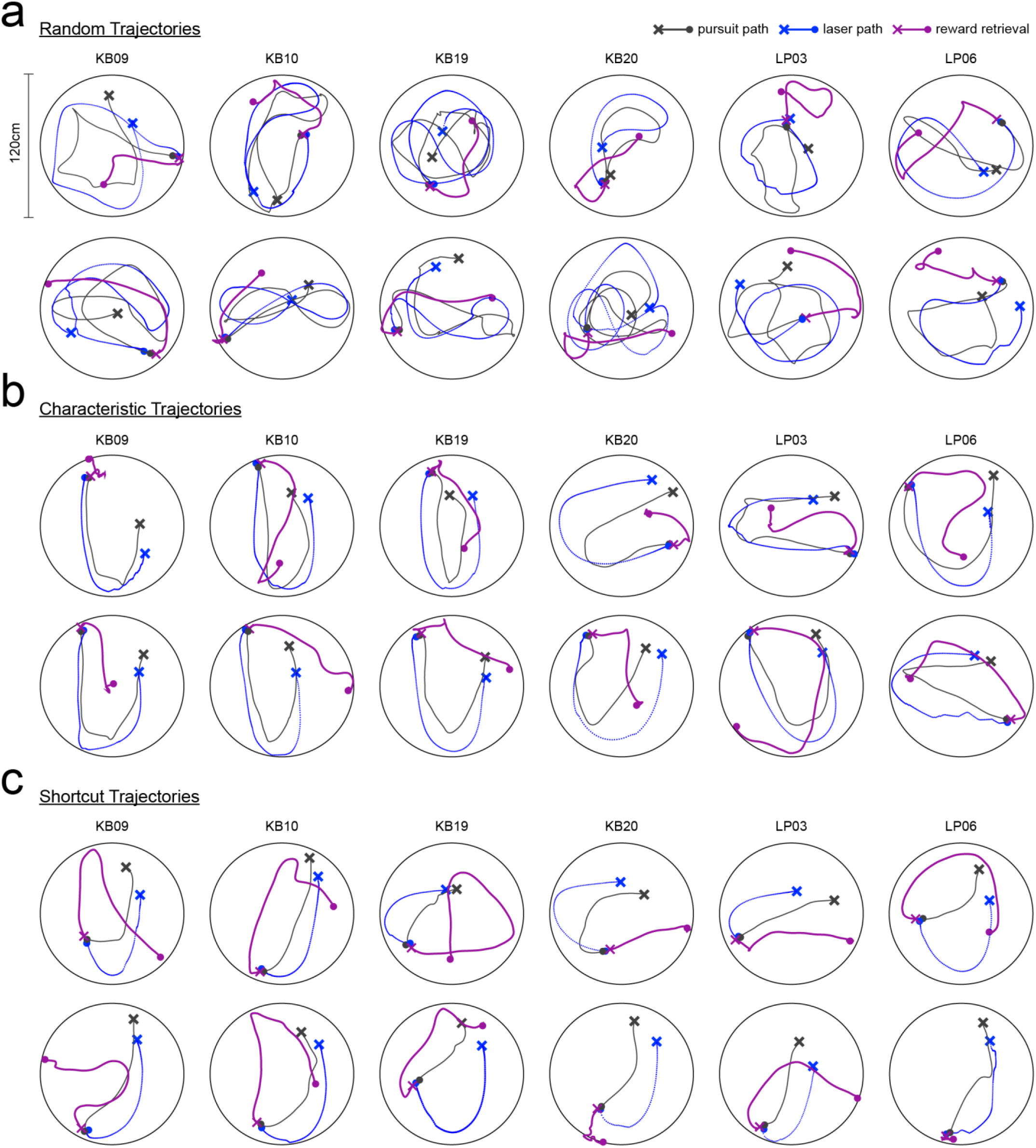
Example random, characteristic, and shortcut trajectories. **a.** 12 example random trajectories. 2 examples are shown for each animal in the study (columns). The trajectory of the rat (dark gray) and target (blue) are shown during chase epochs. Path to reward retrieval is shown in purple. X, start of path. Filled colored circle, end of trial. **b-c.** Same as above but for characteristic and shortcut trajectories, respectively.

**Supplemental Figure 2.**
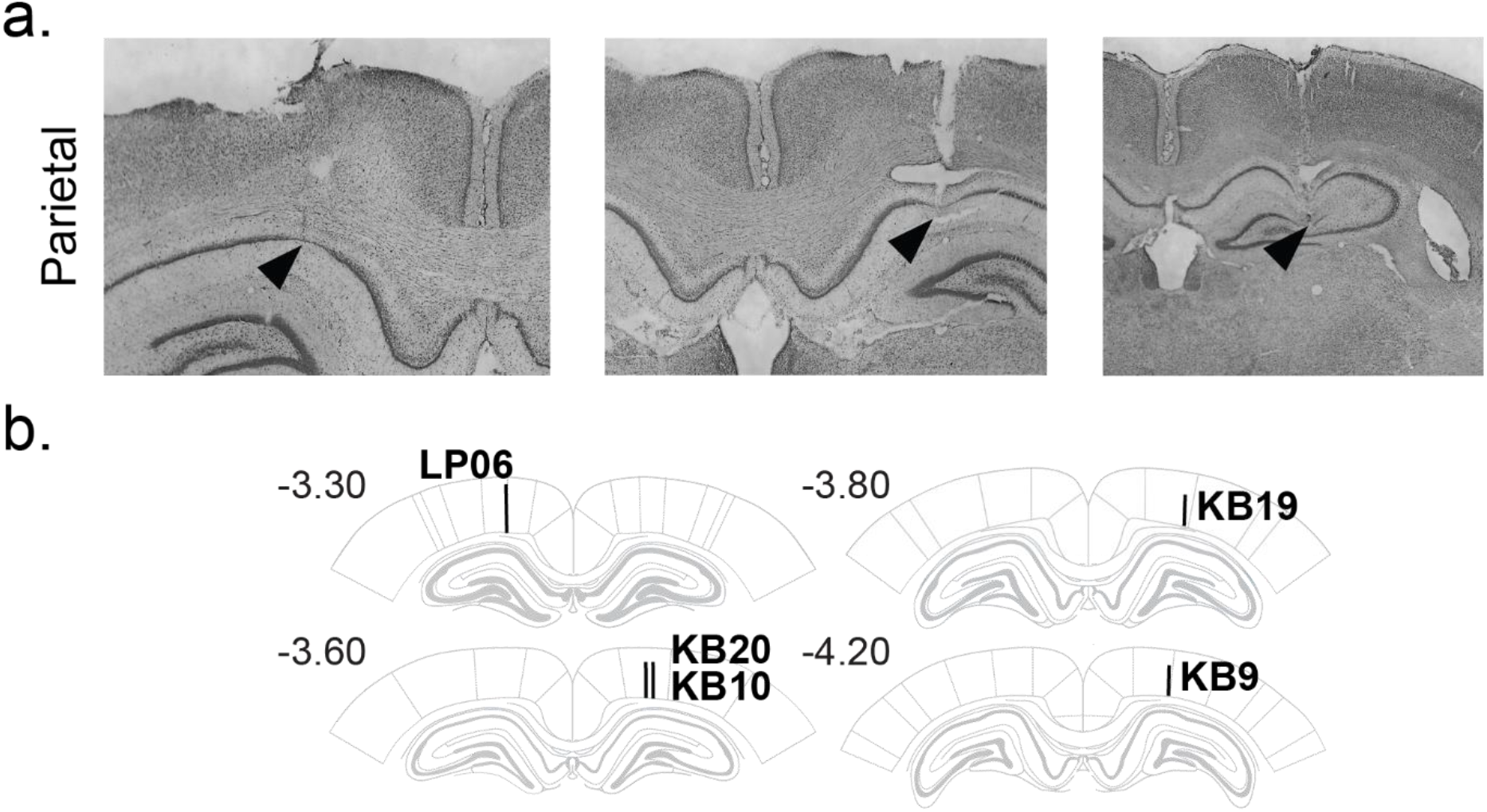
Histological identification of tetrode tracts in posterior parietal cortex. **a.** Black triangles indicate final depth of 3 example tetrode bundles that passed through posterior parietal cortex. Recordings in HPC were not included in analyses. **b.** Schematic of final tetrode tracts for PPC recordings (black).

**Supplemental Figure 3.**
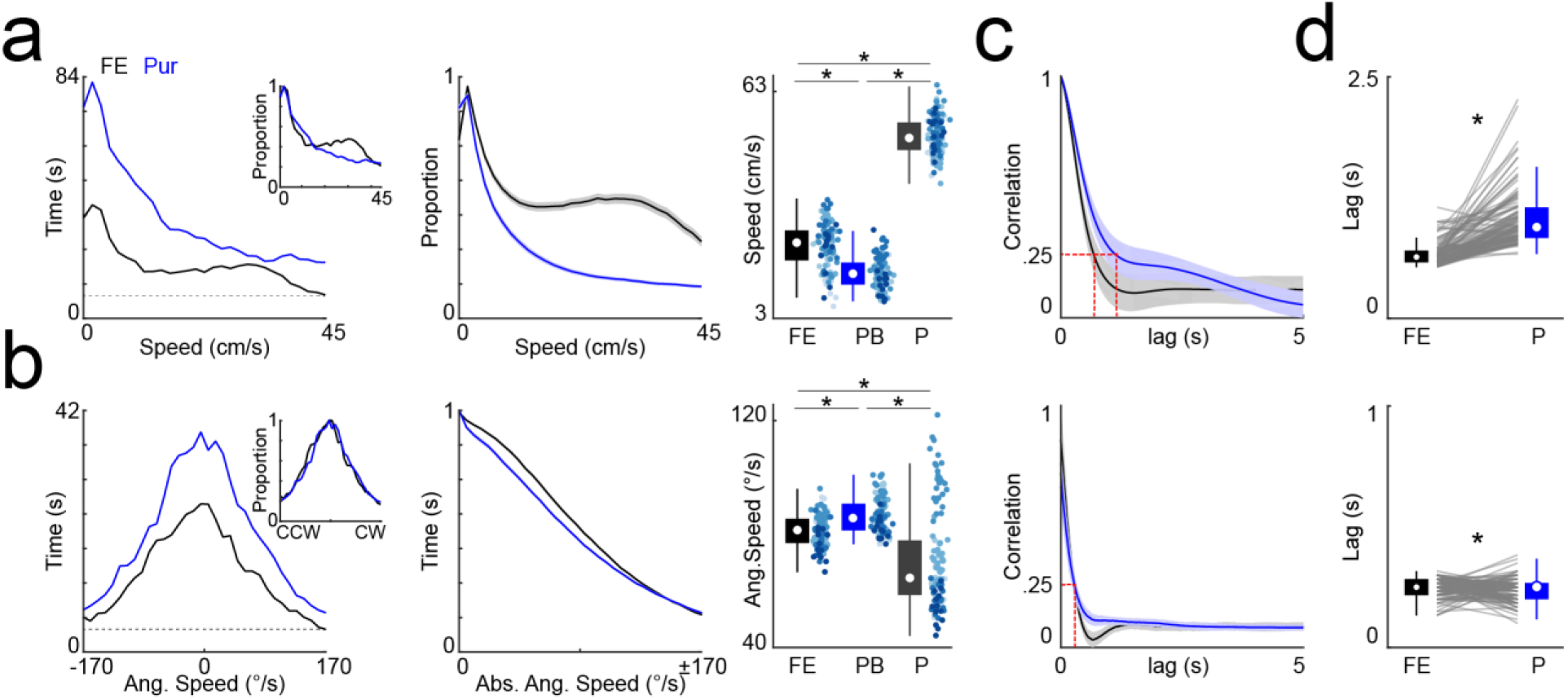
Self-motion differences between target pursuit and free exploration. **a.** Left, histograms of linear speed in pursuit (P, blue) and free exploration (FE, black) sessions on a single day for an individual animal. Absolute time spent at each bin is greater for the P session because of the extended duration when compared to free exploration. Inset shows same data time normalized which highlights different profiles of linear speed amongst P and FE. Horizontal gray dashed line indicates the linear speed with the fewest samples. Construction of behaviorally matched tuning curves for P and FE meant repeated sub-sampling of linear speeds to match the amount of time at the horizontal gray dashed line (minus an additional 1 second). Middle, mean and standard error of normalized histograms for linear speed in P and FE across all animals and sessions. Right, comparison of median linear speed for free exploration (FE), the full pursuit block including pursuits, time between pursuits, and reward consumption (PB), and timeframes in which the animal was actually engaged in a pursuit (P). Colored dots indicate different animals. **b.** Same as in **a**, but for angular speed. **c.** Top, mean and standard deviation of autocorrelation of linear speed across FE and P blocks for all animals. Horizontal red dashed line indicates correlation of 0.25 and intersection of vertical dashed lines with x-axis indicate the temporal lag at which the correlation of linear speed drops below this value. Rightward shift of this intersection during P when compared to free explore indicates that linear speed was correlated for an extended temporal duration during this session when compared to FE. Bottom, same as above but for angular velocity. **d.** Top, comparison of temporal lags wherein autocorrelation of linear speed drops below the 0.25 threshold (red vertical lines in c) for FE and P. Temporal lags in pursuit are significantly greater than in FE, indicating that linear speed was held consistent for more elongated periods of time during chase behavior (n = 113 sessions, median lag FE = 633ms, IQR = 583.3 – 700ms, median lag P = 950ms, IQR = 833 – 1150ms, Wilcoxon rank sum test, z = −11.2, p = 5.52×10^−29^). Below, same comparison as above but for temporal lags of angular speed which was slightly lower in P when compared to FE (n = 113 sessions, median lag FE =250ms, IQR = 216 – 283ms, median lag P = 250ms, IQR = 200 – 267ms, Wilcoxon rank sum test, z = 2.43, p = 0.02).

**Supplemental Figure 4.**
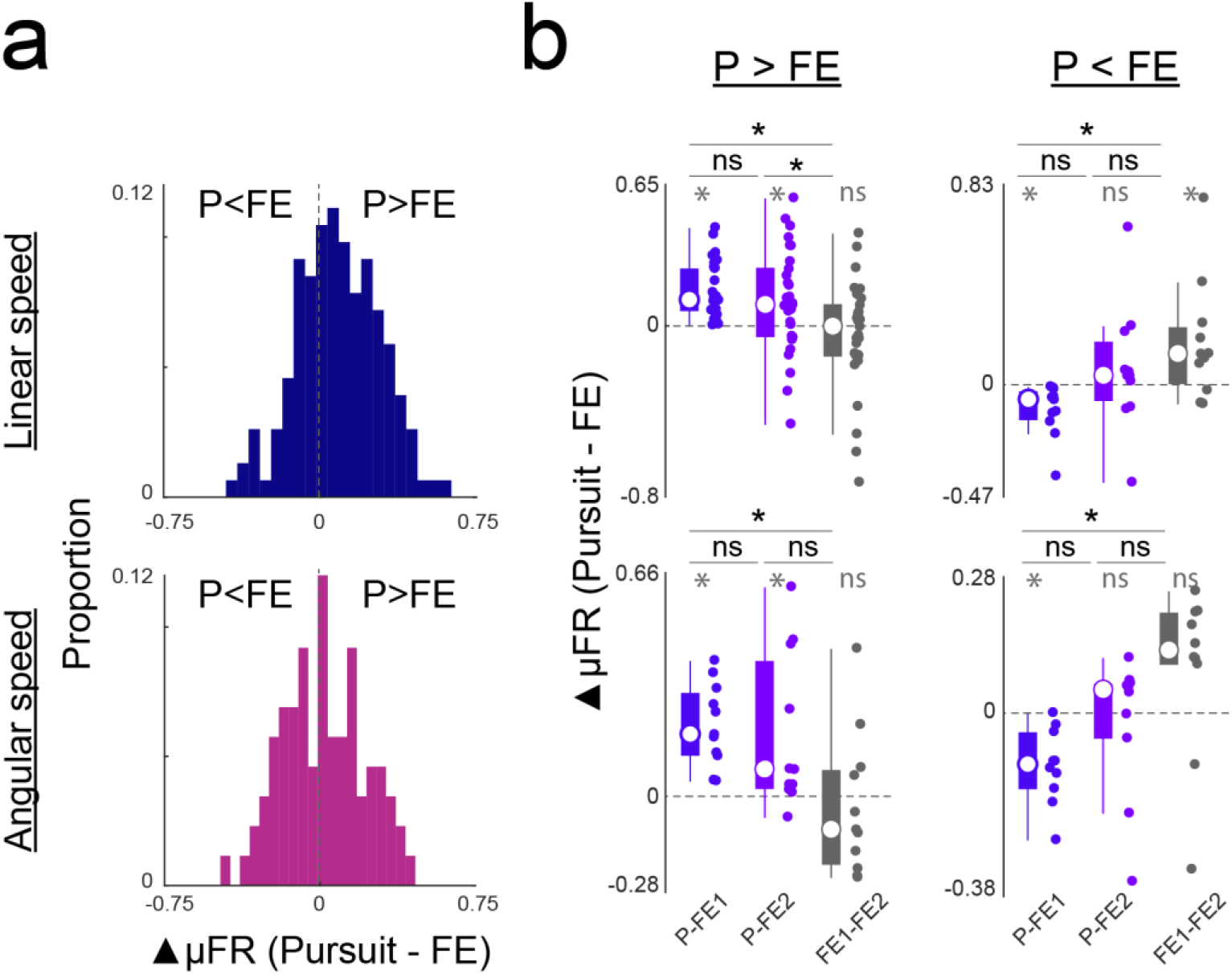
Quantification of mean firing rate differences between pursuit and free exploration. **a.** Top, histogram of mean firing rate differences of linear speed tuning curves that were peak normalized by each neuron’s maximum firing rate across all free explore (FE) and target pursuit sessions. Bottom, same as above but for angular speed tuning curves. Firing rate differences (▲μFR) were computed by subtracting the FE rates from pursuit rates. Accordingly, values to the right of the gray dashed line at zero indicate neurons with greater speed related activity in pursuit than FE. Histograms only depict values for PPC neurons that tested as significantly linear or angular speed sensitive in either FE or pursuit. **b.** Mean firing rate differences between pursuit and FE for the subset of PPC neurons that were significantly self-motion sensitive and recorded in an FE1 – Pursuit – FE2 experimental configuration. Top left, boxplots and single neuron datapoints for mean firing rate differences of linear speed tuning for PPC neurons that had greater activity during pursuit than FE (P > FE). From left to right, boxplots and raw datapoints correspond to firing rate differences between pursuit vs. FE1 (P-FE1), pursuit vs. FE2 (P-FE2), and FE vs FE2 (FE1-FE2). In the left and middle boxplots, firing rate in FE was subtracted from firing rate in Pursuit, thus values above zero indicate greater activity in pursuit. The right boxplot shows firing rate differences when the activity in the second free explore session (FE2) was subtracted from the first FE (FE1). Significance notation in gray indicates rate differences that were significantly non-zero (Signed-rank test for zero median, p < 0.05). Significance notation in black indicates tests between groups (Kruskal-Wallis test, p < 0.05). The top left plot demonstrates that linear speed sensitive PPC neurons composing the P > FE group were systematically modulated by task phase, as this subset had significantly greater activity in pursuit when compared to both FE sessions, while there were no differences in firing rate between FE sessions. Top right, same as top left but for linear speed sensitive PPC neurons with greater activity in FE than pursuit (P < FE). PPC neurons with P < FE have less systematic profiles of firing rate differences across conditions, instead showing more similarity in rate differences for neighboring sessions. Bottom row, same as top row but for differences in angular speed tuning.

**Supplemental Figure 5.**
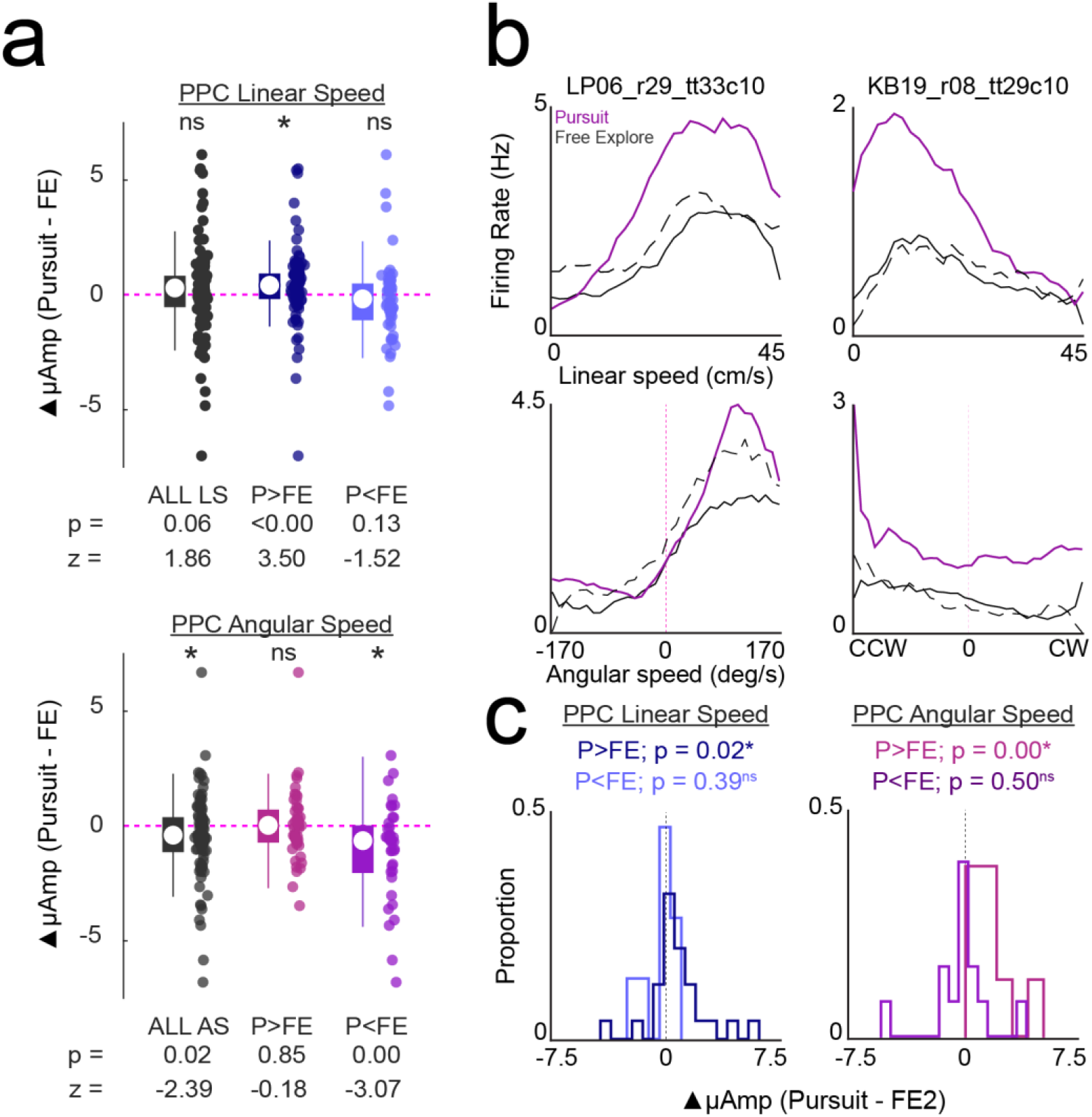
Multiplicative gain modulation of self-motion receptive fields. **a.** Change in amplitude of model fits to linear (top) and angular (bottom) speed tuning curves between pursuit and free exploration sessions. Gray dots and boxplots correspond to all linear (LS) or angular speed (AS) cells. Colored dots and corresponding boxplots show the difference in model amplitude between sessions for PPC neurons with greater mean activity in pursuit (P>FE, dark blue/pink) or free exploration (P<FE, light blue/purple). Values above and below the zero line indicate greater amplitude in pursuit and free explore, respectively. For linear speed sensitive PPC neurons, the population of neurons with greater mean activity in pursuit had significantly greater amplitudes in pursuit than in free explore (Wilcoxon sign rank test for 0 median, p-values and z-values indicated below the plot for each test). For angular speed sensitive PPC neurons, the population of neurons with greater mean activity in free explore had significantly greater amplitudes in free exploration than pursuit. **b.** 2 examples of PPC neurons showing systematic modulation of self-motion receptive field amplitude between pursuit (pink) and both free explore sessions (FE1, solid; FE2, dashed) for linear speed (top). Angular speed tuning curves are shown for the same neurons below. **c.** Quantification of consistent receptive field amplitude changes between pursuit and free explore sessions for PPC neurons. For each region and type of self-motion sensitivity, the subset of neurons with increased (P>FE) and decreased (P<FE) amplitude in pursuit relative to the first free explore were identified. Histograms depict the difference in receptive field amplitude for those populations between pursuit and the second free explore session. If amplitude differences were consistent between pursuit and FE1 and FE2, P>FE neurons would have values skewed positively and P<FE subsets would have values skewed negatively. First plot, PPC neurons with linear speed sensitivity that have greater receptive field amplitudes in pursuit than the first free explore have significantly greater amplitudes between pursuit and the second free explore session (Wilcoxon sign rank for zero median, P>FE n = 25, z = 2.38, p = 0.02), but the P<FE did not have consistent amplitude modulation (Wilcoxon sign rank for zero median, P>FE n = 15, signed rank = 44, p = 0.39). Second plot, PPC neurons with angular speed sensitivity that have greater receptive field amplitudes in pursuit than the first free explore have significantly greater amplitudes between pursuit and the second free explore session (Wilcoxon sign rank for zero median, P>FE n = 8, signed rank = 36, p = 0.01), but the P<FE did not have consistent amplitude modulation (Wilcoxon sign rank for 0 median, P>FE n = 13, signed rank = 35, p = 0.50).

**Supplemental Figure 6.**
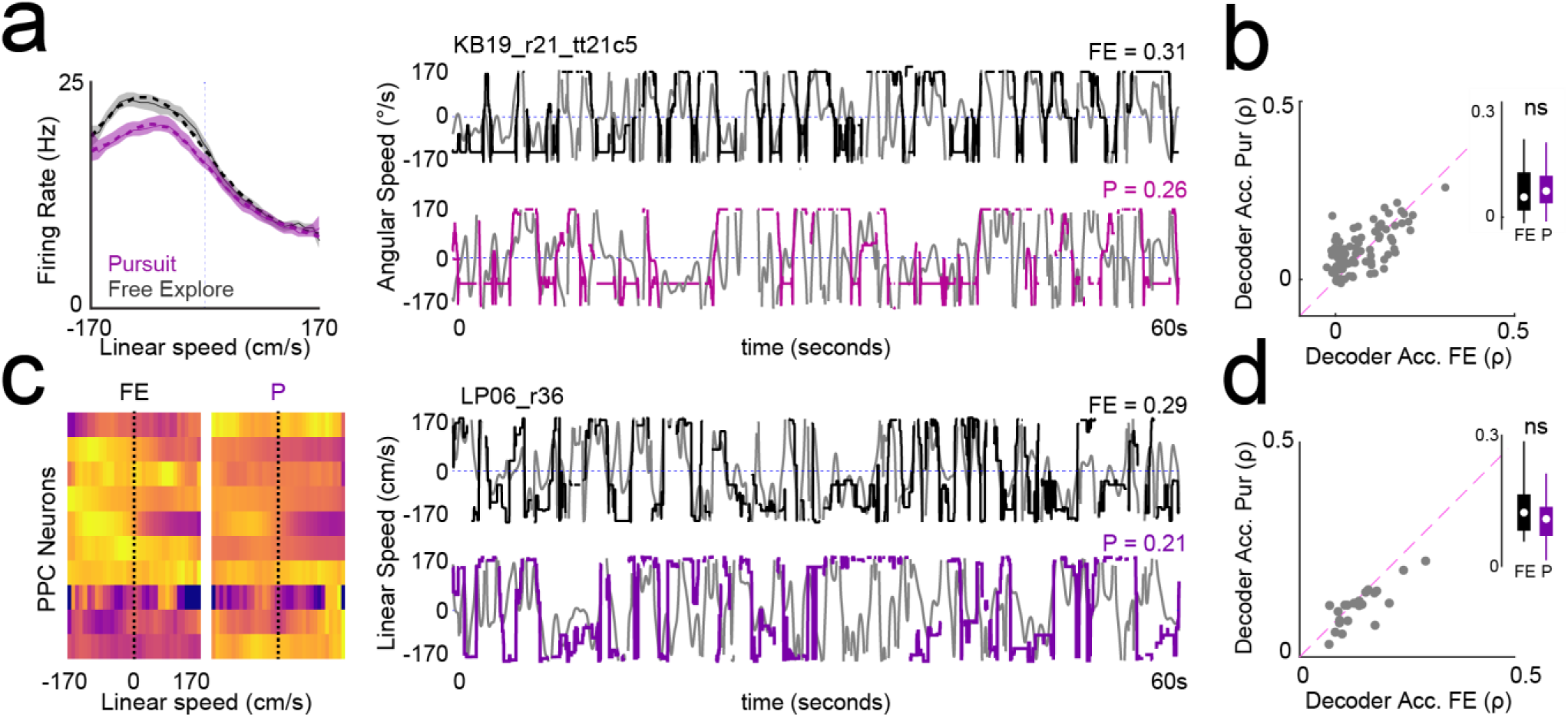
Angular speed decoding accuracy is equivalent between pursuit and free exploration. **a**. Left, example angular speed sensitive neuron with multiplicative gain modulation that decreases rate during pursuit. Right, decoding of angular speed using spiking activity of the same neuron in FE (top) and pursuit (bottom). Decoding accuracy was assessed by correlating the predicted speed (black/pink) with the true speed (gray) and is indicated above each plot. **b.** Angular speed decoder accuracy (Spearman’s rho) is similar for pursuit and FE. Inset, median and IQR of decoder accuracy for FE (black) and pursuit (purple). **c.** Left plot, linearized angular speed tuning curves for 10 simultaneously recorded PPC neurons in FE and P. Rows correspond to the angular speed tuning curves of the same neurons recorded in both conditions and the colormap indicates low (purple) to maximum firing (yellow), peak normalized across all FE and pursuit sessions. Right, corresponding decoder output using this ensemble for FE (top) and pursuit (bottom). Decoder accuracy is indicated above each plot. **d.** Ensemble decoding of angular speed is not significantly different between tasks.

**Supplemental Figure 7.**
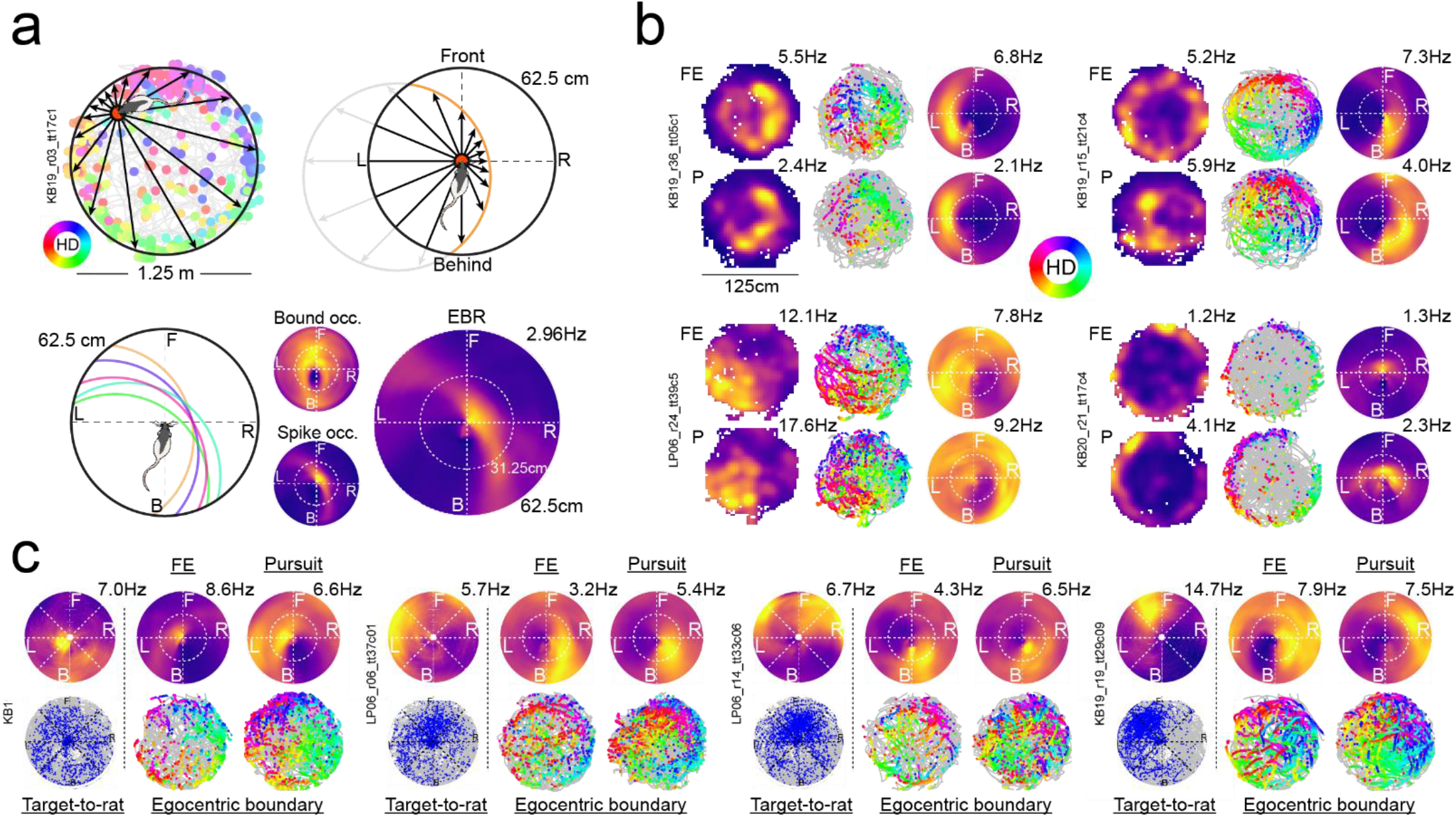
Egocentric boundary vector cells of posterior parietal cortex. **a.** Schematic depicting process for creating egocentric boundary ratemaps (EBR). Top left, in background, trajectory plot with locations of spikes for a single neuron color coded by the rat’s head direction according to the legend on left. For a single spike in the foreground (enlarged in orange), the distance to boundaries in 360° around the animal are determined (a subset shown for clarity). The angle and distance to these boundaries are re-oriented relative to the animal’s head direction at the time of the spike and added to a spike occupancy polar plot with the animal positioned at the origin and facing upwards (top right). Boundaries farther than halfway across the arena (62.5cm) are excluded from the mapping (faded gray). The same process is repeated for all spikes (bottom left) as well as all position samples throughout the session to generate boundary occupancy and spike boundary occupancy maps in egocentric coordinates (bottom insets in middle). EBRs are generated by normalizing the spike occupancy map by the boundary occupancy map (bottom right). This PPC neuron is activated when any environmental boundary is to the right of the animal and slightly ahead of it. **b.** Activation of 4 PPC EBVCs for both the free exploration (top) and pursuit (bottom) sessions. Left columns, two-dimensional ratemaps. Middle columns, trajectory plots with animal trajectory in gray and spikes indicated with colored circles. Colors indicate the head direction of the animal at the time of the spike according to the legend in the middle. Right column, egocentric boundary ratemaps for both sessions. Peak activation indicated above all allocentric and egocentric ratemaps. **c.** Activation of 4 PPC EBVCs that were simultaneously sensitive to the egocentric position of the visual target relative to the animal. Left column, rat-to-target egocentric ratemaps and corresponding trajectory plots. Rat-to-target egocentric ratemaps constructed in a similar manner as EBRs but instead reflect activation as a function of the egocentric position of the visual target relative to the animal. Trajectory plots show every position of the target (within 40cm), relative to the animal, in gray. Target positions at the time of spikes are depicted in blue dots. Right two columns, corresponding egocentric boundary ratemaps and trajectory plots for the same neuron in free exploration and pursuit show that, in addition to mapping the position of the visual target, these neurons simultaneously map the position of boundaries.

## Supplemental Videos

**sVideo 1. An example pseudorandom trajectory in a pre-surgery rat in the light.**

**sVideo 2. An example pseudorandom trajectory in a pre-surgery rat in the light.**

**sVideo 3. An example pseudorandom trajectory in a post-surgery rat in the dark.**

**sVideo 4. An example characteristic trajectory #1 in a pre-surgery rat in the light.**

**sVideo 5. An example characteristic trajectory #2 in a pre-surgery rat in the light.**

**sVideo 6. An example characteristic trajectory #2 in a post-surgery rat in the dark.**

**sVideo 7. An example shortcut on characteristic trajectory #1 in a pre-surgery rat in the light.**

**sVideo 8. An example shortcut on characteristic trajectory #1 in a pre-surgery rat in the light.**

**sVideo 9. An example shortcut on characteristic trajectory #2 in a post-surgery rat in the dark.**

